# Gene regulatory networks controlling differentiation, survival, and diversification of hypothalamic Lhx6-expressing GABAergic neurons

**DOI:** 10.1101/2020.05.21.106963

**Authors:** Dong Won Kim, Kai Liu, Zoe Qianyi Wang, Yi Stephanie Zhang, Abhijith Bathini, Matthew P Brown, Sonia Hao Lin, Parris Whitney Washington, Changyu Sun, Susan Lindtner, Bora Lee, Hong Wang, Tomomi Shimogori, John L.R. Rubenstein, Seth Blackshaw

**Affiliations:** Solomon H. Snyder Department of Neuroscience, Johns Hopkins University School of Medicine, Baltimore, MD, 21205, USA; Department of Ophthalmology, Johns Hopkins University School of Medicine, Baltimore, MD, 21205, USA; Department of Neurology, Johns Hopkins University School of Medicine, Baltimore, MD, 21205, USA; Center for Human Systems Biology, Johns Hopkins University School of Medicine, Baltimore, MD, 21205, USA; Institute for Cell Engineering, Johns Hopkins University School of Medicine, Baltimore, MD, 21205, USA; Kavli Neuroscience Discovery Institute, Johns Hopkins University School of Medicine, Baltimore, MD, 21205, USA; Nina Ireland Laboratory of Developmental Neurobiology, Department of Psychiatry, UCSF Weill Institute for Neurosciences, University of California, San Francisco, San Francisco, CA 94158, USA; Center for Neuroscience, Korea Institute of Science and Technology (KIST), Seoul, 02792, Korea; RIKEN Center for Brain Science, Laboratory for Molecular Mechanisms of Brain Development, 2-1 Hirosawa, Wako, Saitama, 351-0198, Japan; Genentech, South San Francisco, CA 94080

## Abstract

GABAergic neurons of the hypothalamus regulate many innate behaviors, but little is known about the mechanisms that control their development. We previously identified hypothalamic neurons that express the LIM homeodomain transcription factor Lhx6, a master regulator of cortical interneuron development, as sleep-promoting. In contrast to telencephalic interneurons, hypothalamic Lhx6 neurons do not undergo long-distance tangential migration and do not express cortical interneuronal markers such as *Pvalb*. Here, we show that *Lhx6* is necessary for the survival of hypothalamic neurons. *Dlx1/2*, *Nkx2-2*, and *Nkx2-1* are each required for specification of spatially distinct subsets of hypothalamic Lhx6 neurons, and that Nkx2-2+/Lhx6+ neurons of the zona incerta are responsive to sleep pressure. We further identify multiple neuropeptides that are enriched in spatially segregated subsets of hypothalamic Lhx6 neurons, and that are distinct from those seen in cortical neurons. These findings identify common and divergent molecular mechanisms by which Lhx6 controls the development of GABAergic neurons in the hypothalamus.

## Introduction

Although much is now known about both the diversity and development of GABAergic neurons of the telencephalon^1,2^, far less is known about their counterparts in the hypothalamus, where over 20% of neurons are GABAergic^3^. Previous work shows that hypothalamic GABAergic neuronal precursors first appear in a domain that separates the anterodorsal and posteroventral halves of the developing hypothalamus, and is delineated by expression of transcription factors that regulate the development of telencephalic GABAergic neurons, including *Dlx1/2* and *Arx* ^4–8^. Within this structure, which has been termed the intrahypothalamic diagonal/tuberomammillary terminal (ID/TT), nested expression domains of LIM homeodomain family genes are observed, in which expression of *Lhx1*, *Lhx8*, and *Lhx6* delineates the anterior-posterior axis of the ID/TT^4^. *Lhx1* is essential for the terminal differentiation and function of neurons in the master circadian oscillator in the suprachiasmatic nucleus^9–11^. Lhx6-expressing neurons in the zona incerta (ZI) of the hypothalamus are sleep-promoting and activated by elevated sleep pressure, and hypothalamic-specific loss of function of *Lhx6* disrupts sleep homeostasis^12^.

Lhx6 has been extensively studied in the developing telencephalon. It is essential for the specification, migration, and maturation of GABAergic neurons of the telencephalon – particularly the cortex and hippocampus^13,14^. Lhx6 is expressed in the medial ganglionic eminence (MGE) of the embryonic telencephalon (Fig. 1A, B), where it is coexpressed with both Nkx2-1 and Dlx1/2^15–17^. Shh induces expression of *Nkx2-1*^18^, which in turn directly activates *Lhx6* expression^16,19^. Nkx2-1, in turn, cooperates with Lhx6 to directly activate the expression of multiple other genes that control cortical interneuron specification and differentiation, including *Sox6* and *Gsx2* ^20,21^. Furthermore, *Lhx6* is both necessary and sufficient for the tangential migration of the great majority of interneuron precursors from the MGE to their final destinations in the cortex and hippocampus^17,22,23^. Finally, Lhx6 expression persists in mature interneurons that express parvalbumin (Pvalb) and somatostatin (Sst), and is necessary for their expression^22^.

**Figure 1.**
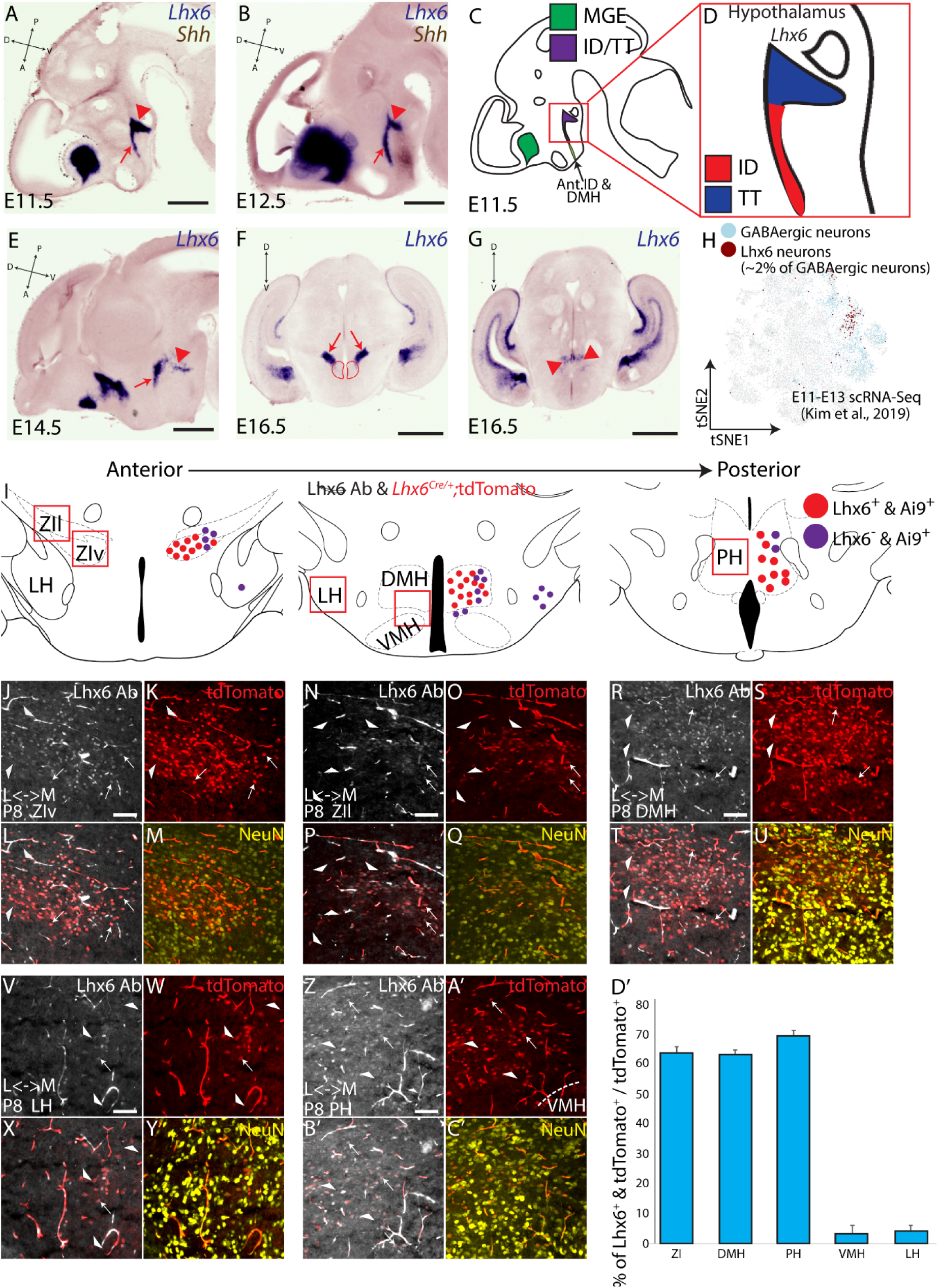
Distribution of hypothalamus *Lhx6*-expressing neurons. (A, B, E-G) *In situ* hybridization showing *Lhx6* (blue) and *Shh* (brown) at E11.5 (A), E12.5 (B), E14.5 (E), shown in sagittal planes, and E16.5 (F, G) in coronal planes. Red arrows (A, B, E) indicate the intrahypothalamic diagonal (ID), red arrowheads (A, B, E) indicate the tuberomammillary terminal (TT), black arrows (D) indicate migrated telencephalic *Lhx6*-expressing cells (tangential migration from the medial ganglionic eminence to the cortex). Red arrows in F indicate the ZI and red circle in F indicates the DMH, and red arrows in G indicate the PH. (C, D) Schematics showing the distribution of telencephalic (green) and hypothalamic (purple) *Lhx6*-expressing cells at E11.5, with ID (red) and TT (blue) are highlighted in D (E11.5) Note anterior domains to the ID that shows a weak and transient *Lhx6* expression during development (Ant.ID = anterior ID, DMH = dorsomedial hypothalamus). (H) scRNA-Seq from E11-E13 hypothalamus scRNA-Seq from^24^ showing the distribution of neurons that express GABAergic markers (blue, *Dlx1/2*, *Gad1/2*, and *Slc32a1*) and Lhx6-expressing GABAergic neurons (brown) that are ~2% of all hypothalamic GABAergic neurons during development. (I) Schematic distribution of Lhx6-expressing neurons across the dorsolateral hypothalamus (red = neurons that continue to express *Lhx6*, purple = neurons that transiently expressed *Lhx6*) across ZIv (zona incerta ventral), ZIl (zona incerta lateral), LH (lateral hypothalamus), DMH (dorsomedial hypothalamus), VMH (ventromedial hypothalamus), PH (posterior hypothalamus). (J-C’) Lhx6 antibody staining (grey), tdTomato expression from *Lhx6*^*Cre/+;*^;*Ai9* line (red), NeuN (yellow) in ZIv (J-M), ZIl (N-Q), DMH (R-U), LH (V-Y), PH (Z-C’). L = lateral, M = medial. White arrowheads show neurons that transiently expressed Lhx6, and white arrows show neurons continue to express Lhx6). (D’) A bar graph showing the percentage of tdTomato+ and Lhx6-expressing neurons over the total number of tdTomato^+^ neurons. Scale bar = 0.45 mm (A), 0.5 mm (B, F, G), 0.55 mm (E), 0.1 mm (I-C’).

The functional role of Lhx6 in hypothalamic development has not been previously investigated. However, previous studies imply this may differ in certain key ways from its function in the developing telencephalon. Notably, the hypothalamic domain of Lhx6 expression only partially overlaps with that of *Nkx2-1*^4^. Furthermore, in sharp contrast to cortical interneurons, Lhx6 is not co-expressed with either Pvalb or Sst has been reported in the ZI^12^. In this study, we sought to determine the extent to which gene regulatory networks controlling the development of hypothalamic Lhx6 neurons diverge from those that control the development of telencephalic Lhx6 neurons. We find that hypothalamic Lhx6 regulates neuronal differentiation and survival. Further, we observe extensive molecular heterogeneity among mature hypothalamic Lhx6 neurons and a lack of overlap with annotated subtypes of Lhx6-expressing cortical interneurons. Combinatorial patterns of transcription factor expression delineate spatial subdomains of Lhx6 expression within the ID/TT, and we find that *Nkx2-1*, *Nkx2-2*, and *Dlx1/2* each regulate expression of Lhx6 in largely non-overlapping domains. Finally, Lhx6 neurons derived from *Nkx2-2*-expressing precursors are activated by sleep pressure. These findings identify mechanisms by which Lhx6 can regulate the development of hypothalamic GABAergic neurons, and more broadly, how diverse subtypes of hypothalamic neurons can be generated during development.

## Results

### Distribution of hypothalamic Lhx6-expressing neurons

Our previous work has indicated that Lhx6 is expressed in two continuous yet distinct domains of the developing hypothalamus: the intrahypothalamic diagonal (ID) and the more posterior tuberomammillary terminal (TT)^4,5^. We next sought to more carefully determine the expression pattern of *Lhx6* and its putative regulators during early hypothalamic development. High-quality chromogenic *in situ* hybridization (ISH) detects both the ID and TT domain of *Lhx6* expression at E11.5, E12.5, and E14.5 (Fig. 1A-E). By E16.5, hypothalamic *Lhx6*-expressing neurons are observed in the zona incerta (ZI) and dorsomedial hypothalamus (DMH), in a pattern that broadly corresponds to the earlier ID domain, while expression in the posterior hypothalamus (PH) in turn broadly corresponds to the TT domain (Fig. 1F, G). This closely matches the pattern of hypothalamic *Lhx6* expression previously reported in adults^12^. Lhx6-expressing neurons are only a small minority of hypothalamic GABAergic neurons^12^, with scRNA-Seq revealing that only ~2% of all hypothalamic GABAergic neuronal precursors (defined by *Gad1/2* and *Dlx1/2* expression) express *Lhx6* between E11 and E13 (Fig. 1H)^24^.

This regional pattern of hypothalamic *Lhx6* expression is broadly similar to that reported for *Lhx6*^*Cre/+*^;*Ai9* mice (Fig. 1J-D’)^12^, with ~65-70% of tdTomato-expressing neurons in the ZI, DMH, and PH of *Lhx6*^*Cre/+*^;*Ai9* postnatal mice also continuing to express Lhx6 (Fig. 1J-U, D’). Notably, we see a few tdTomato-expressing neurons in other hypothalamic regions, with the largest numbers found in adjacent structures such as the VMH (surrounding VMH) and lateral hypothalamus (LH), although only ~5% of these tdTomato-expressing neurons still express Lhx6 (Fig. 1V-D’). This shows that, in contrast to telencephalic interneuron precursors, hypothalamic Lhx6 cells do not appear to undergo long-distance tangential migration, and that hypothalamic Lhx6-expressing cells that do undergo short-range tangential dispersal during early development generally repress Lhx6 expression as they mature.

### *Lhx6* is necessary for the survival of hypothalamic neurons

These findings led us to investigate other potential differences in the Lhx6 function in hypothalamic neurons relative to the telencephalon. While Lhx6 does not regulate the survival of cortical interneuron precursors^22^, hypothalamic-specific loss of function of *Lhx6* leads to substantial changes in sleep patterns^12^, raising the possibility that Lhx6 may be necessary for the viability or proper functions of these neurons.

To investigate this possibility, we tested P8 *Lhx6*^*CreER/CreER*^ mice, in which a CreER cassette has been inserted in frame with the start codon to generate a null mutant of *Lhx6*, to determine if read-through transcription of endogenous *Lhx6* could be detected in the hypothalamus (Fig. 2A). Chromogenic *in situ* hybridization of telencephalic structures such as the amygdala and cortex reveals that *Lhx6*-expressing cells are still detected in both regions, although the number of *Lhx6*-expressing cells in the cortex is substantially reduced in *Lhx6*^*CreER/CreER*^ mice relative to *Lhx6*^*CreER/+*^ heterozygous controls (Fig. 2B-M). This is consistent with the severe reduction in tangential migration of cortical interneurons seen in *Lhx6*-deficient mice^22,23^. In the hypothalamus, however, no read-through transcription of *Lhx6* was detected (Fig. 2J, M). This implies that, in contrast to its role in the telencephalon where *Lhx6* is necessary for the tangential migration and proper laminar positioning^21,23^, hypothalamic *Lhx6* is required to promote neuronal survival and/or to activate its expression.

**Figure 2.**
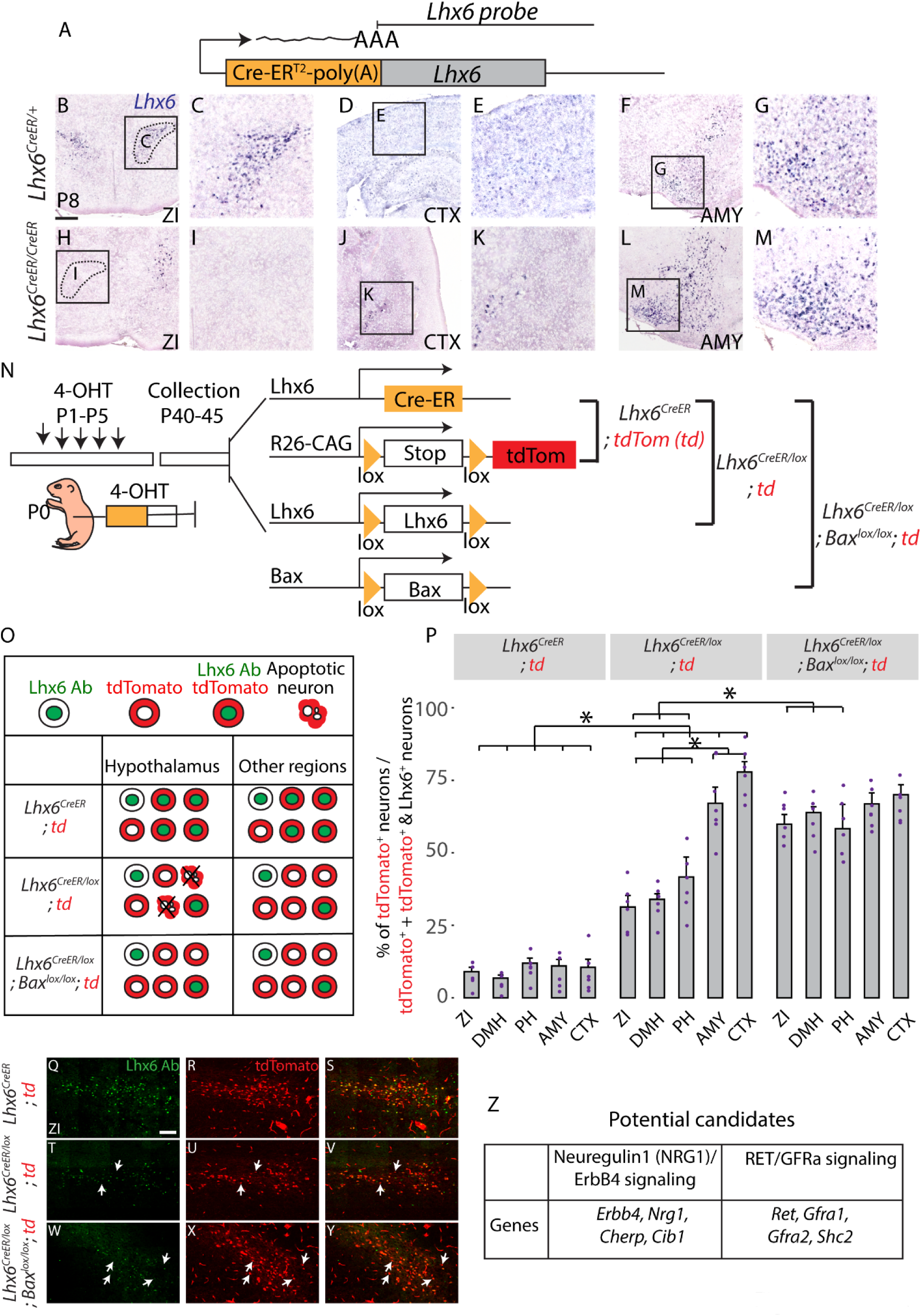
Lhx6 in the hypothalamus is necessary for neuronal survival. (A) *Lhx6*^*CreER*^ knock-in site (JAX #010776) and location of the *Lhx6* probe used to detect read-through transcription. (B-M) Coronal planes showing *Lhx6* mRNA expression in the zona incerta (ZI, B, C, H, I), amygdala (AMY, F, G, L, M), and cortex (CTX, D, E, J, K) in control (*Lhx6*^*CreER/+*^, B-G) and mutant (*Lhx6*^*CreER/CreER*^, H-M) at P8. Note *Lhx6* mRNA is not detected in the Lhx6-deficient hypothalamus but is detected in the telencephalon. (N) Schematic diagram showing the overall design of the experiment from 3 genotypes (1. *Lhx6*^*CreER/+*^;*Ai9*, 2. *Lhx6*^*CreER/lox*^;*Ai9*, 3. *Lhx6*^*CreER/lox*^;;*Bax*^*lox/lox*^;*Ai9*). (O) Schematic diagram showing the overall outcome of the experiment. (P) A bar graph showing the percentage of tdTomato^+^ / (tdTomato^+^ & tdTomato^+^/Lhx6^+^) across 3 genotypes in 5 different brain regions. * indicates *p < 0.05*. DMH = dorsomedial hypothalamus, PH = posterior hypothalamus. (Q-Y) Representative images of 3 genotypes (1. *Lhx6*^*CreER/+*^;*Ai9* (Q-S), 2. *Lhx6*^*CreER/lox*^;*Ai9* (T-V), 3. *Lhx6*^*CreER/lox*^;*Bax*^*lox/lox*^;*Ai9* (W-Y) in ZI. More images of different brain regions are available in Fig. S1. (Z) Potential candidate genes from bulk RNA-Seq (Fig. S2) controlling cell survival can be regulated by *Lhx6* in hypothalamic Lhx6^+^ neurons: Neuregulin-ErbB4 signaling and Gdnf signaling pathways (Fig. S3, S4). Scale bar = 0.6 mm (B-M), 100 μm (Q-Y).

To distinguish between these possibilities, we sought to determine whether neonatal loss of function of *Lhx6* would lead to the death of Lhx6-expressing neurons. This was done using the genetic fate mapping of *Lhx6*-deficient neurons. Using a series of 4-Hydroxytamoxifen (4-OHT) injections between P1 and P5 in *Lhx6*^*CreER/+*^*Ai9* and *Lhx6*^*CreER/lox*^;*Ai9* mice, we labeled *Lhx6*-expressing cells with tdTomato while also simultaneously disrupting *Lhx6* function in a subset of Lhx6-expressing neurons in *Lhx6*^*CreER/lox*^ mice (Fig. 2N). We then quantified the number of neurons that expressed both tdTomato and Lhx6 protein at P45, as well as the number of neurons that only expressed tdTomato. Expression of the only tdTomato indicates that a cell has lost expression of Lhx6, either as a result of Cre-dependent disruption of the *Lhx6* locus or as a result of normal repression of expression during postnatal development (Fig. 2O). In both hypothalamic and telencephalic regions in *Lhx6*^*CreER/+*^;*Ai9* mice, we observed that the fraction of neurons that only express tdTomato was only 10-15% of the number of neurons expressing both Lhx6 and tdTomato (Fig. 2P-S, Fig. S1). This indicates that the great majority of neurons in both regions that express *Lhx6* in neonates continue to do so at P45. However, when we performed this same analysis in *Lhx6*^*CreER/lox*^;*Ai9* mice, we found that while 75% of tdTomato-expressing neurons in the cortex and amygdala remain even in the absence of detectable Lhx6 protein, a substantially smaller fraction of tdTomato-expressing neurons are detected in the absence of Lhx6 protein in the ZI, DMH, and PH (Fig. 2P, T-V, Fig. S1).

This is consistent with *Lhx6* playing a selective role in regulating the survival of Lhx6-expressing hypothalamic neurons. To directly address this hypothesis, we next generated *Lhx6*^*CreER/lox*^;*Bax*^*lox/lox*^;*Ai9* mice, with loss of function of *Bax* predicted to selectively prevent apoptosis in Lhx6-expressing neurons^25^. When Cre recombinase activity was induced using the same protocol, we observed that the fraction of tdTomato-expressing neurons that lacked Lhx6 expression was indistinguishable from that seen in cortex and amygdala (Fig. 2P, W-Y, Fig. S1).

These data indicate that Lhx6 is selectively required for the survival of hypothalamic Lhx6-expressing neurons. To determine whether *Lhx6* is also required for normal differentiation of these cells, we next conducted RNA-Seq analysis on sorted tdTomato-expressing hypothalamic cells from P10 *Lhx6*^*CreER/+*^;*Ai9* and *Lhx6*^*CreER/lox*^;*Bax*^*lox/lox*^;*Ai9* mice (Fig. 2Z, Fig. S2). We observe that *Lhx6*^*CreER/lox*^;*Bax*^*lox/lox*^;*Ai9* mice show no change in expression of markers of GABAergic neurons, including *Gad1*, *Gad2*, *Slc32a1*. However, substantially increased expression of genes expressed in mitotic neural progenitors, including *Ccna1*, *Aurka, Msx1*, and *Msx2* (Fig. S2, Table S1), is observed, along with a decreased expression of axon guidance/growth factors such as *Sema3c*, *Sema4d*, *Sema5a*. Notably, we also observe ectopic expression of genes that are not normally found in the brain but are expressed in germline stem cells (*Sycp1*), testes (*Ccdc144b, Samd15, Stag3*) mucosa (*Slc12a8*), colon (*Nlrp6*), liver (*Tfr2*), heart (*Popdc2, Spta1*), and cochlear hair cells (*Pdzd7*)^26^. This suggests that, as in telencephalic neurons, Lhx6 is not required for expression of GABAergic markers^22,23,21,23^, but might be required to repress inappropriate expression of genes expressed both in neural progenitor and in non-neuronal cells. This does not, however, exclude the possibility that these may be in part induced as a result of the loss of function of *Bax*.

Genetic and biochemical analyses have identified several genes as direct or indirect Lhx6 targets in the developing telencephalon^18,19,21,23,27^. These include *Shh*, the transcription factors *Arx*, *Cux2*, *Mafb*, *Nkx2-1*; as well as *Sst* and chemokine receptors such as *Cxcr4*, *Cxcr7*, and *Erbb4*. To identify genes and signaling pathways that are strong candidates for selectively regulating survival of hypothalamic Lhx6 neurons, the bulk RNA-Seq data from P10 *Lhx6*^*CreER/+*^;*Ai9* neurons were directly compared to profiles obtained from FACS-isolated *Lhx6-GFP* positive and negative hypothalamic and cortical neurons that were collected at E15.5, P8 (Fig. 2Z, Fig. S2), since regulation of hypothalamic Lhx6 in cell survival is evident during embryonic and early neonatal periods and we expected to detect potential signaling pathways at both datasets. Genes found to be enriched in hypothalamic samples of bulk RNA-Seq data were then compared to single-cell RNA-Sequencing (scRNA-Seq) datasets of hypothalamic Lhx6-expressing neurons collected at E15.5 and P8 (Fig. 2Z, Fig. S2)^24^, and a core set of Lhx6-regulated genes that were selectively enriched in hypothalamic Lhx6-expressing neurons was thus identified.

We observe that many previously identified Lhx6 targets either show little detectable expression in wildtype hypothalamic Lhx6 neurons, (*Cux2*, *Mafb*, *Sst*, *Cxcr4/7*) or else showed no detectable change in expression following *Lhx6* loss of function (*Arx*, *Nkx2-1*). One notable exception is the Neuregulin receptor *Erbb4*, which has been shown to be necessary for tangential migration and differentiation of MGE-derived immature Lhx6-expressing cortical interneurons^28–30^. *Erbb4* is both highly expressed in hypothalamic Lhx6 neurons, and its expression is strongly *Lhx6*-dependent (Fig. 2Z, Fig. S2). Since Neuregulin signaling is also neurotrophic in many cell types^31^, this suggested that the loss of neuregulin signaling could be a potential mechanism behind the apoptotic death of *Lhx6*-deficient hypothalamic cells. Indeed, we observed that additional components of the both the Neuregulin (*Nrg1*) and Gdnf (*Ret, Gfra1, Gfra2*) neurotrophic signaling pathways were selectively enriched in hypothalamic Lhx6 neurons (Fig. 2Z, Fig. S2), a finding which was confirmed using fluorescent *in situ* hybridization and scRNA-Seq (Fig. S3).

### Diverse subtypes of Lhx6-expressing neurons are found in the postnatal hypothalamus

Our previous work^12^ showed that adult ZI Lhx6-expressing neurons do not highly express traditional markers of MGE Lhx6^+^ derived GABAergic neurons of the cortex. No ZI Lhx6-expressing neurons co-express Pvalb and Sst, with only a small subset expressing Npy^12^. We thus hypothesized that subtypes of Lhx6 neurons in the postnatal hypothalamus might be diverged substantially from those present in the cortex^32^, and might be more molecularly heterogeneous.

scRNA-Seq analysis of P8 Lhx6-eGFP neurons from the hypothalamus that expressed high levels of Lhx6 mRNA shows that these neurons express a diverse pool of neuropeptides and neurotransmitters that are not expressed in telencephalic Lhx6-expressing neurons, including *Gal*, and *Trh* (Fig. 3, Fig. S4, Table S2). Other markers that are specific to distinct subsets of cortical Lhx6 neurons were expressed in hypothalamic Lhx6 neurons, such as *Pnoc*, *Tac1, Nos1*, and *Th*. Hypothalamic Lhx6-expressing neurons do not express *Pvalb*, but a small fraction expresses *Npy* and *Cck*. We also identified a rare subpopulation of hypothalamic Lhx6-expressing neurons in the PH that co-express *Sst*, although these are absent in more anterior regions (Fig. 3, Fig. S4). *Tac1* is expressed broadly in cortical and hypothalamic Lhx6-expressing neurons. Similar patterns of gene expression are observed in scRNA-Seq data obtained from Lhx6 neurons in the adult hypothalamus of mice that are older than P30 (Fig. S5)^24,33,34^. However, all these enriched markers (neuropeptides and neurotransmitters) are not specific to Lhx6-expressing neurons but rather expressed broadly in hypothalamic GABAergic neurons across nuclei (Fig. S6).

**Figure 3.**
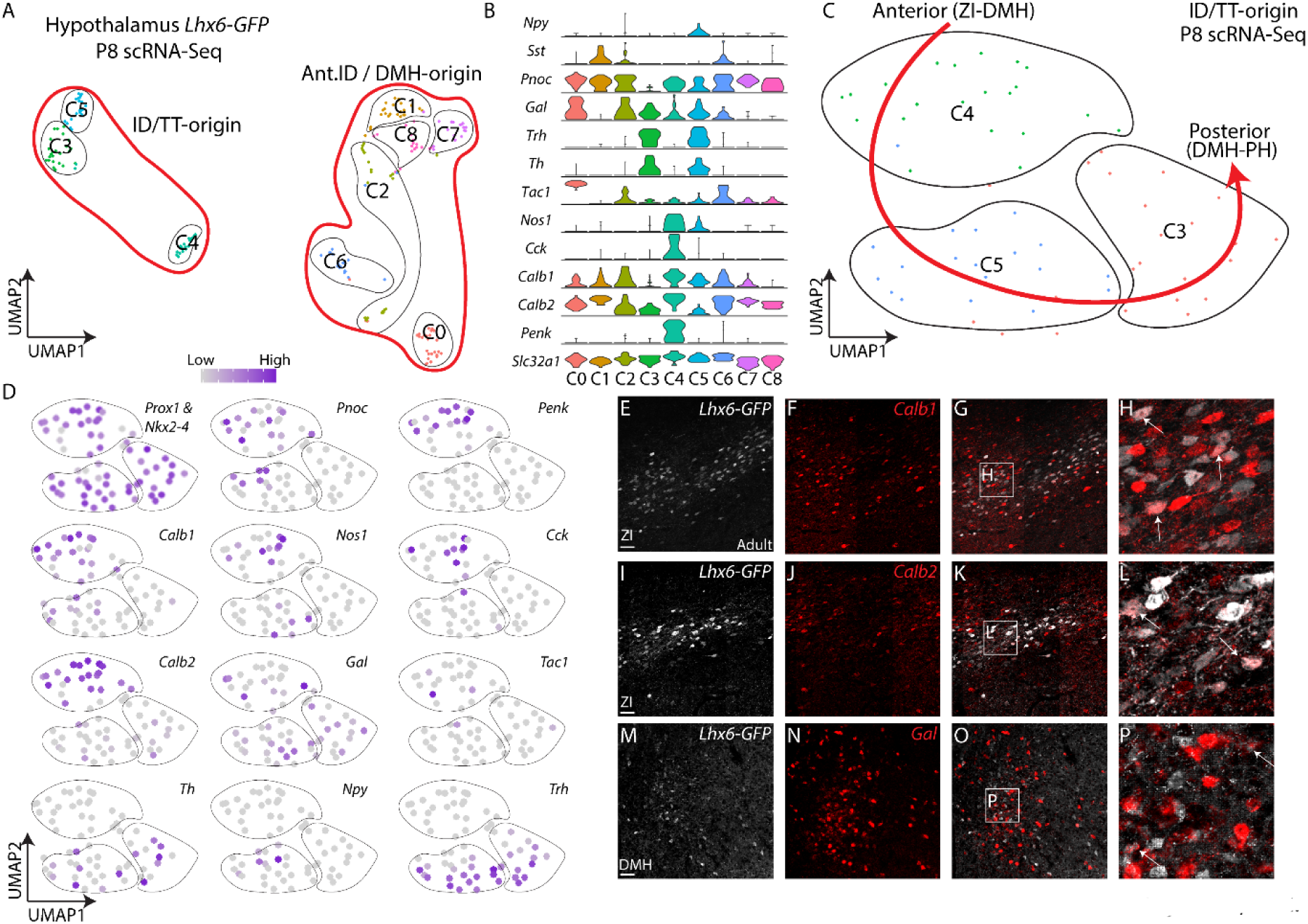
Diverse subtypes of mature hypothalamic Lhx6-expressing neurons. (A) UMAP plot showing hypothalamic Lhx6-expressing GABAergic neurons at P8 scRNA-Seq. (B) Violin plots showing key markers in individual clusters in A. (C) UMAP plot showing Lhx6-expressing neurons originated from ID and TT. (D) UMAP plots showing the distribution of diverse neuropeptides and neurotransmitters across ID and TT derived Lhx6-expressing neurons. (E-P) Fluorescent *in situ* hybridization showing *Lhx6-GFP* (grey) with *Calb1* (red, E-H), *Calb2* (red, I-L), and *Gal* (red, M-P). Scale bar = 50 μm.

Mature Lhx6 hypothalamic neurons were organized into three major clusters that showed close similarity to the two subdomains of the ID and the main TT region observed at E12.5, and in turn appear to represent individual subtypes of Lhx6 neurons that are differentially distributed along the anteroposterior axis of the hypothalamus, and which may correspond to Lhx6 neurons of the ZI, DMH, and PH, respectively (Fig. 3, Fig. S4). Lhx6 neurons express a mixture of *Pnoc*, *Penk*, *Calb1*, *Calb2*, *Cck* in both the ZI and DMH, whereas *Tac1* is more restricted to the ZI. *Npy* and *Nos1* are enriched in DMH Lhx6 neurons. *Th*, *Trh*, *Gal* are located in the region spanning the DMH and PH, while *Sst* is expressed only in a small subset of PH Lhx6 neurons.

### scRNA-Seq identifies molecular markers of spatially distinct domains of hypothalamic Lhx6 neurons

Lhx6-expressing neurons of the postnatal hypothalamus are molecularly diverse and distributed across a broad region of the dorsolateral hypothalamus. We hypothesized that this diversity is regulated by multiple transcription factors that control the specification of region-specific subtypes of Lhx6-expressing neurons.

To identify these anatomically and molecularly distinct Lhx6-expressing domains in the hypothalamus, we performed single-cell RNA sequencing (scRNA-Seq) with the *Lhx6-GFP* line at E12.5 and E15.5. At E12.5 and E15.5, scRNA-Seq analysis readily distinguishes the ID, TT, and hinge domains (Fig. 4a, Fig. S7, S8). By E12.5, all Lhx6 cells in the hypothalamus express the early neuronal precursor marker *Dcx*, as well as the synaptic GABA transporter *Slc32a1*, but do not express progenitor markers (e.g. *Fabp7* and *Ascl1*), indicating these Lhx6-expressing cells are postmitotic neurons as early as E12.5 (Fig. 4B, C). In addition, weak *Lhx6* expression was observed in *Lhx1* and *Lhx8* co-expressing neurons of the anterior ID cluster, which are *Nkx2-1*^+^ (Fig. 4D, E), and give rise to GABAergic neurons in the suprachiasmatic nucleus and DMH, although little or no *Lhx6* mRNA was detected in these neurons after E13.5 (Fig. 1, Fig. 6)^4,10^. We observed that *Dlx1/2*, *Nkx2-2*, and *Nkx2-1* are differentially expressed in the ID, hinge, and TT domains, respectively, at both ages (Fig. 4D, E, Fig. S8). These 3 transcription factors are each shown as putative regulons of the ID, hinge, and TT domains respectively (Fig. 4F). Furthermore, we observe several molecularly distinct cell clusters that have not been previously described. The first cluster expresses low levels of *Nkx2-1*, but high levels of *Prox1* and *Sp9*, transcription factors that are highly expressed in the developing prethalamus. This may therefore correspond to a dorsal subdomain of the TT located adjacent to the hinge domain (Fig. 4D, E). We also observe a distinction between more proximal and distal domains of the ID, based on the expression of *Nefl, Dlx6, Nefm, Lhx1*, and *Nr2f1*.

**Figure 4.**
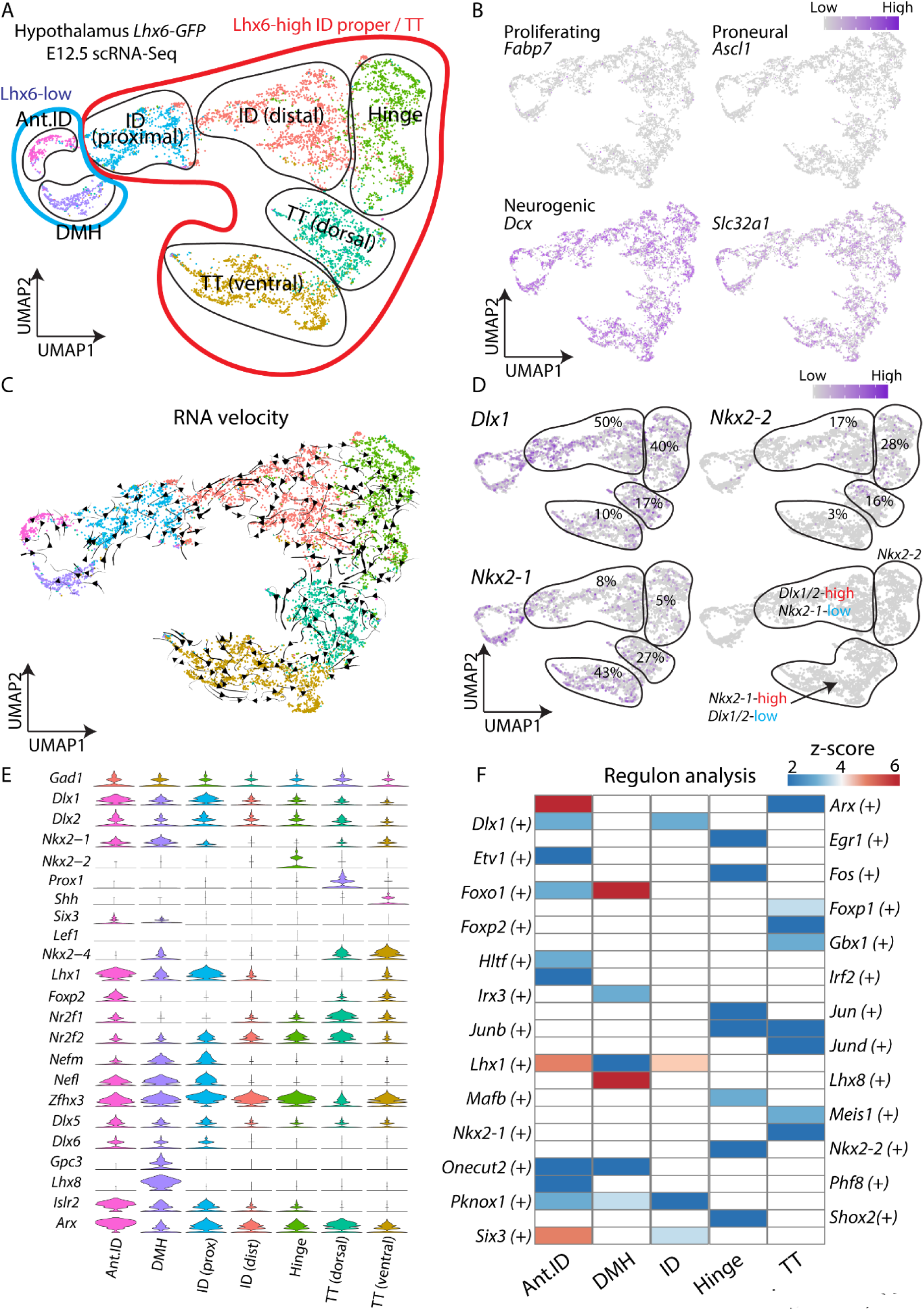
scRNA-Seq identifies molecular markers of spatially distinct domains of hypothalamic Lhx6 neurons. (A) UMAP plot showing different Lhx6-expressing hypothalamic regions at E12.5. (B) UMAP plots showing a lack of expression of *Fabp7* (proliferating cells), *Ascl1* (proneural), whereas the neuronal precursor marker *Dcx* and the GABAergic neuronal marker *Slc32a1* is highly expressed. (C) UMAP plot with RNA velocity trajectories. (D) UMAP plots showing expression and percentage of ID, hinge, and TT Lhx6 neurons expressing *Dlx1/2*, *Nkx2-2*, and *Nkx2-1*. (E) Violin plots showing expression of key transcription factors (and other genes) that are highly expressed in individual domains. (F) A heatmap showing z-scores of significantly differentially expressed regulons among Lhx6^+^ hypothalamic regions. Note regulon activity of *Dlx1* in the ID, *Nkx2-2* in the hinge, and *Nkx2-1* in the TT. Ant.ID = anterior ID, DMH = dorsomedial hypothalamus.

In all, five molecularly distinct clusters of neurons that strongly express Lhx6 could be resolved in the embryonic hypothalamus (Fig. 4). These can be distinguished not only by the expression of different subsets of transcription factors at E12.5, but also by more conventional markers of cell identity such as neuropeptides and calcium-binding proteins such as *Sst, Tac1, Pnoc, Islr2, Gal*, and *Npy* at E15.5 (Fig. S8, Table. S3, S4). We also observed clusters that were located in the hinge and TT region at E12.5 (Fig. S8 cluster 4 and 7), but which postnatally expressed markers that are restricted to neurons at the most anterior domain of hypothalamic Lhx6 neurons. These markers include *Nfix*, *Nfib*, and *Tcf4* (Fig. S8, Fig. 3, Table. S2-S4).

These molecularly distinct domains of hypothalamic Lhx6 neurons were also visualized using traditional two-color ISH with *Nkx2-1, Nkx2-2, Arx*, and *Prox1* probes (Fig. S9). This also confirms that *Shh* is only expressed in dorsal TT Lhx6 neurons, while *Six3* is expressed only in the weakly Lhx6-expressing neurons in the anterior ID. scRNA-Seq showed that *Lef1*, which is expressed broadly in the ID and TT region at E12.5, was expressed in only very few Lhx6 neurons at both E12.5 and E15.5 (Fig. 4, Fig. S9), indicating that *Lef1* and *Lhx6* are not extensively co-expressed.

### *Dlx1/2*, *Nkx2-2*, and *Nkx2-1* mediate patterning of discrete spatial domains of hypothalamic Lhx6 neurons

*Dlx1/2, Nkx2-2*, and *Nkx2-1* are selectively expressed in the ID, hinge, and TT domains, respectively. Since these three transcription factors were also identified as putative regulons from scRNA-Seq analysis, we sought to investigate their function in regulating *Lhx6* expression in more detail. Using *Lhx6-GFP* mice, which faithfully recapitulate the endogenous expression pattern of *Lhx6*^12^, we integrated bulk RNA-Seq analysis obtained at E15.5 and P0 from hypothalamus with age-matched ATAC-Seq data to cross-reference our scRNA-Seq result (Fig. 5A). We further sought to investigate similarities and differences in gene expression and chromatin accessibility in age-matched hypothalamic and telencephalic Lhx6-expressing neurons (Fig. 5), since the role of Lhx6 in development of telencephalic interneurons is extensively studied, and it is therefore critically important to connect these findings to prior work characterizing Lhx6 mechanisms of action in forebrain development.

**Figure 5.**
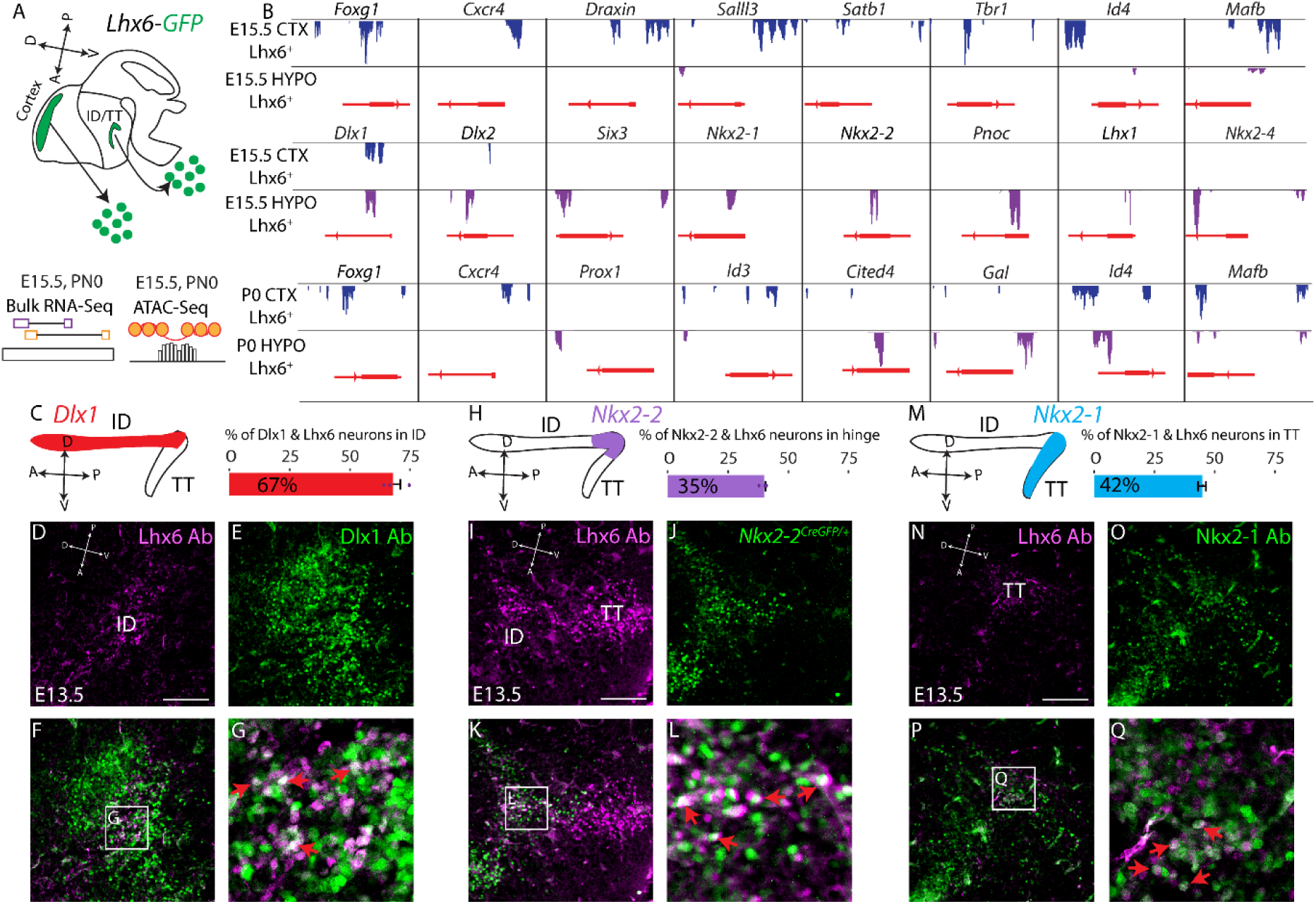
*Dlx1/2, Nkx2-2*, and *Nkx2-1* are expressed in distinct spatial domains of hypothalamic Lhx6 neurons. (A) Schematic showing bulk RNA-Seq and bulk ATAC-Seq pipelines from flow-sorted *Lhx6-GFP+* neurons of the cortex and hypothalamus at E15.5 and P0. (B) Peaks in ATAC-Seq footprinting sites showing potential transcription factor binding sites near the promoter regions of differentially expressed genes from bulk RNA-Seq data in the cortex and hypothalamus at E15.5 and P0. (C) Schematic showing the ID/TT of the developing hypothalamus and expression of *Dlx1/2* (left), and the percentage of ID Lhx6-expressing neurons that co-express *Dlx1* (right). (D-G) Immunostaining with Lhx6 (purple) and Dlx1 (green) of E13.5 hypothalamus, showing co-localization of Lhx6 and Dlx1 in the ID of the hypothalamus (G, red arrows). (H) Schematic showing ID/TT of the developing hypothalamus and expression of *Nkx2-2* (left) and the percentage of hinge Lhx6-expressing neurons that co-express *Nkx2-2* (right). (I-L) Immunostaining with Lhx6 (purple) and GFP in *Nkx2-2*^*CreGFP/+*^ (green) of E13.5 hypothalamus show co-localization of Lhx6 and *Nkx2-2GFP* between the ID and TT (hinge region) of the hypothalamus (L, red arrows). (M) Schematic showing the ID/TT of the developing hypothalamus and expression of *Nkx2-1* (left) and the percentage TT Lhx6-expressing neurons that co-express *Nkx2-1* (right). (N-Q) Immunostaining with Lhx6 (purple) and Nkx2-1 (green) of E13.5 hypothalamus, showing co-localization of Lhx6 and Dlx1 in the ID of the hypothalamus (Q, red arrows). Scale bar = 50 μm.

At E15, many region-specific differences in gene expression were observed between hypothalamic and telencephalic Lhx6-expressing neurons, particularly for transcription factors. We observed enriched expression of *Six3*, *Nkx2-2*, and *Nkx2-4* in the hypothalamus. As predicted by earlier studies, we observed enriched expression of the telencephalic marker *Foxg1*, *Satb2*, and *Nr2e1*^*35,36*^, in the cortex (Fig. 5B, Table S5).

However, expression of genes broadly expressed in GABAergic neurons showed no significant differences, including *Nkx2-1*, and *Dlx1/2*. At P0, hypothalamic Lhx6 neurons continued to show enriched expression for multiple transcription factors, including *Prox1*, *Foxp2*, and *Nhlh2*. Hypothalamic Lhx6 neurons show little detectable expression of the cortical interneuron markers *Pvalb, Sst*, and *Npy*, but we observed a higher level of *Gal* and *Pnoc* at P0 in hypothalamic Lhx6 neurons. Relative to *Lhx6*-negative hypothalamic neurons, we also observed a higher level of transcription factors such as *Dlx1*, *Onecut1*, *Pax5*, and *Nkx2-2* in hypothalamic Lhx6 neurons compare to the rest of the hypothalamus at E15.5, as well as a higher level of *Tac1* and *Pnoc* at P0 (Fig. S10, Table S6).

Regions of accessible chromatin identified by ATAC-Seq were, as expected, clustered in the proximal promoter and intronic regions of annotated genes in all samples profiled (Fig. S10, Table S7, S8). Region-specific differences in chromatin accessibility frequently corresponded to differences in mRNA expression. For instance, proximal promoter and/or intronic regions of *Foxg1, Npy, Pvalb*, and *Sst* were selectively accessible in cortical Lhx6 neurons, while those of *Nkx2-2, Sall3*, and *Gal* were accessible only in the hypothalamus at both E15.5 and P0 (Fig. 5B, Table S7, S8, Fig. S10). However, substantial differences in chromatin accessibility were also observed for *Nkx2-1* and *Dlx1/2* at both E15.5 and P0, implying that different gene regulatory networks may control the expression of these genes in hypothalamus and cortex (Table S7, S8).

To determine whether any of the spatial domains of Lhx6 expression could closely resemble telencephalic Lhx6 cells, we compared E12.5 hypothalamic scRNA-Seq results to data previously obtained from E13.5 MGE^37^. These data confirmed that, while transcription factors such as *Nkx2-1*, *Dlx1/2*, and *Lhx8* are broadly expressed in Lhx6 MGE cells, they are not expressed (*Lhx8*) or expressed only in discrete subsets (*Nkx2-1*, and *Dlx1/2*) of hypothalamic Lhx6 neurons. No identified subset of hypothalamic Lhx6 neurons resembled MGE Lhx6 cells (Fig. S11A, B, Table S9).

With substantial differences between hypothalamic and telencephalic Lhx6-expressing neurons in both gene expression and chromatin accessibility, we reasoned that the transcriptional regulatory networks identified as controlling the development of telencephalic Lhx6-expressing neurons would not broadly apply in developing hypothalamus. Thus, based on both scRNA-Seq data and analysis of our ATAC-Seq data, as well as our previous work^4,24^, three previously mentioned transcription factors -- *Nkx2-1*, *Dlx1/2*, and *Nkx2-2* -- emerged as strong candidates for regulating specific domains of hypothalamic Lhx6 neurons. *Nkx2-1* is required for Lhx6 expression in the telencephalon^16,19^ and is expressed in the TT, but not ID, domain in the hypothalamus^4,24^, while *Dlx1/2* are required for tangential migration of cortical interneurons and are also broadly expressed in both cortical and hypothalamic Lhx6 neurons^4,6,8,38^. *Nkx2-2*, in contrast, is expressed only in the hypothalamus in a zone immediately dorsal to the region of *Nkx2-1* expression^4,39^.

Each of these transcription factors is expressed in discrete spatial domains that overlap with distinct subsets of hypothalamic Lhx6 neurons at E13.5 (Fig. 5C-Q). *Dlx1* was strongly expressed in the ID (Fig. 5C-G, Fig. S11C-H), but not the TT. Nkx2-2, in contrast, selectively demarcated the region joining the ID and TT (Fig. 5H-L), which we have termed the hinge domain. Nkx2-1 was selectively expressed in the TT region, but essentially absent from the ID and hinge domain (Fig. 5M-Q, Fig. S11I-K). These spatial differences in the expression of Dlx1 and Nkx2-1 in hypothalamic Lhx6 neurons are preserved at E17.5, where *Dlx1* is enriched in the more anterior ZI and DMH (Fig. S12), and Nkx2-1 expression is enriched in the PH (Fig. S12A-L). Furthermore, unlike the MGE Lhx6-expressing cells, Dlx1 and Nkx2-1 formed mutually exclusive expression domains in the ID and TT (Fig. S11F-K). However, we observed a much more even distribution of Nkx2-2/Lhx6 neurons across the ZI, DMH, and PH, which could indicate either short-range tangential dispersal of hinge neurons or widespread induction of Nkx2-2 expression in Lhx6 neurons at later ages (Fig. S12M-X). These results indicate that distinct spatial domains of hypothalamic Lhx6 expression can be delineated by combinatorial patterns of homeodomain transcription factor expression.

To determine the final location of *Nkx2-1* expressing Lhx6 neurons, we next used fate-mapping analysis, in which *Nkx2-1*^*CreER/+*^;*Ai9* mice^40^ were labeled with 4-OHT at E11 (Fig. S13). At E18, tdTomato expression was detected in the majority of Lhx6-expressing neurons in the amygdala and cortex (Fig. S13) as expected^16,19^, but we observed anterior-posterior bias in the distribution of tdTomato-expressing neurons in the hypothalamus that closely matched the location of Lhx6/Nkx2-1 expressing neurons at earlier ages. We observe that only a small fraction (~10%) of ZI Lhx6-expressing neurons, which correspond to the most anterior region of Lhx6 expression at later developmental ages^12^, were labeled with tdTomato. In contrast, a much larger fraction of PH Lhx6 neurons, corresponding to the most posterior domain of Lhx6 expression, were tdTomato-positive. This implies that Nkx2-1/Lhx6-expressing neurons of the TT primarily give rise to Lhx6 neurons found in the posterior hypothalamus, but that a small fraction may undergo tangential migration to more anterior structures such as the ZI. This was also shown with immunostaining of Nkx2-1 and Lhx6-expressing neurons at E17.5 (Fig. S12Y-J’).

We next investigated whether loss of function of *Nkx2-1*, *Nkx2-2*, and *Dlx1/2* led to the loss of spatially-restricted hypothalamic expression of Lhx6. We first examined *Nkx2-1*^*CreER/CreER*^ mice, in which targeted insertion of the CreER cassette generates a null mutation in *Nkx2-1*^40^. This leads to severe hypoplasia of the posteroventral hypothalamus, as previously reported for targeted *Nkx2-1* null mutants^41^. The ventrally-extending TT domain of Lhx6 expression is not detected in *Nkx2-1*-deficient mice at E12.5, but the Nkx2-1-negative ID domain persists (Fig. 6, Fig. S14). Fate mapping analysis, in which *Nkx2-1*^*CreER/+*^;*Ai9* and *Nkx2-1*^*CreER/CreER*^;*Ai9* mice were injected with tamoxifen at E11 and analyzed at E18, indicate that surviving Lhx6 neurons in the ID region represent a mixture of tdTomato-positive and -negative neurons, and confirm that a subset of these surviving neurons derived from *Nkx2-1*-expressing precursors. As previously reported, no Lhx6-expressing neurons are detected in the mutant cortex (Fig. S14).

**Figure 6.**
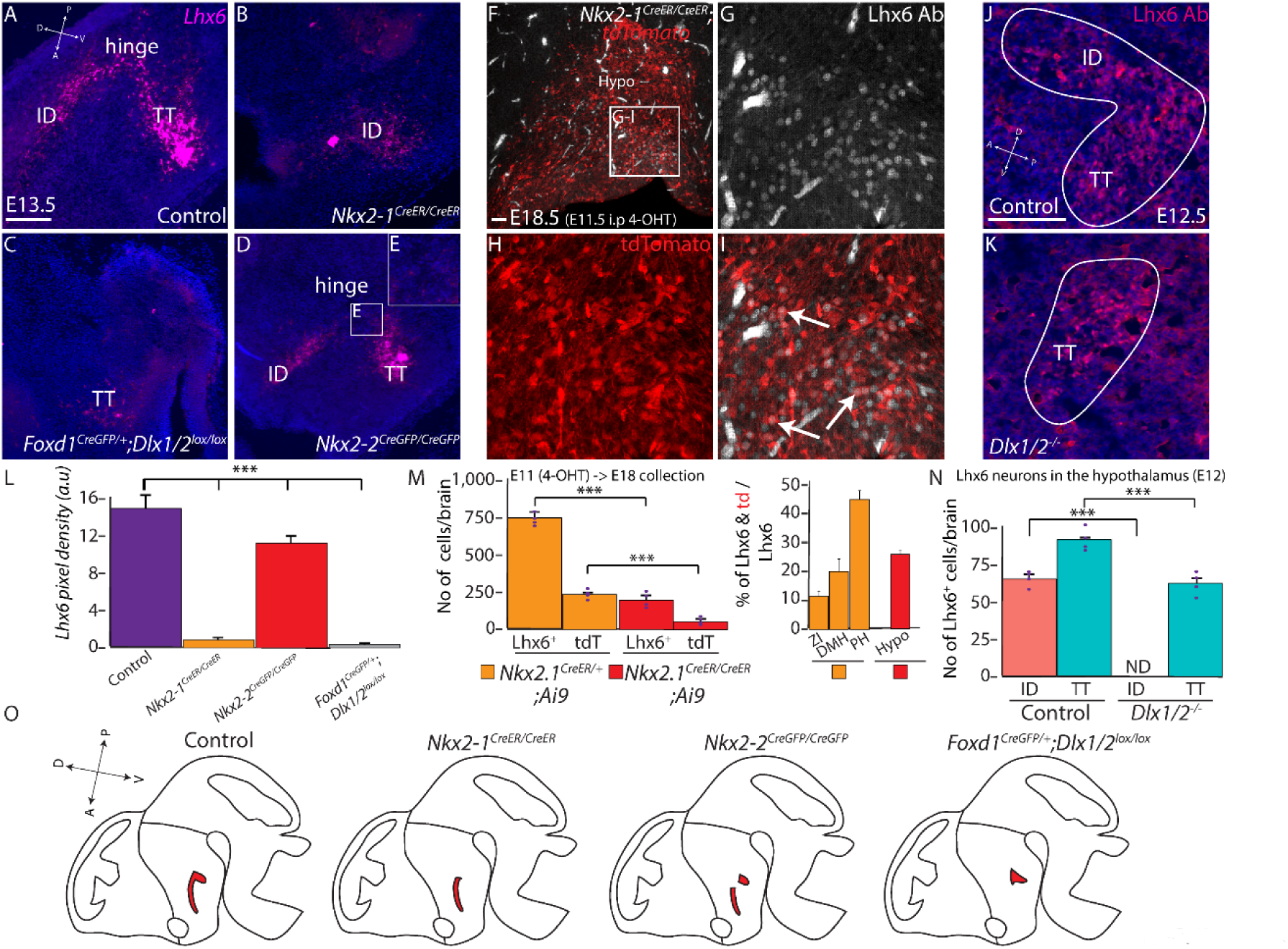
*Dlx1/2, Nkx2-2*, and *Nkx2-1* mediate patterning of discrete spatial domains of hypothalamic Lhx6 neurons. (A-E, L) RNAscope showing *Lhx6* expression (magenta) in control (A), *Nkx2-1*^*CreER/CreER*^ (B), *Foxd1*^*Cre/+*^:*Dlx1/2*^*lox/lox*^ (C), and *Nkx2-2*^*CreGFP/CreGFP*^ (D-E). Pixel density of *Lhx6* is shown across all 4 groups in (L). (F-I, M) *Nkx2-1*^*CreER/CreER*^;*Ai9* (4-OHT treatment at E11.5, collection at E18.5) showing Lhx6 expression (grey) and tdTomato (red) in the hypothalamus. Arrows in I indicate Lhx6^+^ and tdTomato^+^ neurons. Raw number of Lhx6^+^ or tdTomato^+^ (tdT) neurons (left) and percentage of Lhx6^+^ & tdTomato^+^ neurons (right) in *Nkx2-1*^*CreER/+*^;*Ai9* (Fig. S5) and *Nkx2-1*^*CreER/CreER*^/*Ai9*. (J-K, N) Lhx6 expression (magenta) in control and Dlx1/2^−/−^ at E12 ID and TT. The number of Lhx6-expressing neurons in the ID and TT are shown in (N). (O) Schematic diagram showing distribution of *Lhx6* expression across 4 groups. ID = intrahypothalamic diagonal, TT = tuberomamillary terminal. Scale bar = 50 μm. *** *p < 0.05*.

We next generated null mutants of *Nkx2-2* in the same manner, generating mice homozygous for a knock-in CreGFP cassette that disrupts expression of the endogenous *Nkx2-2* locus^42^. In this case, we observe a loss of Lhx6 expression in the hinge region, located between the posterior ID and dorsal TT (Fig. 6, Fig. S14). Finally, we examined the phenotype of mice deficient for *Dlx1/2*, examining both global knockouts^43^ and *Foxd1*^*Cre/+*^;*Dlx1/2*^*lox/lox*^ mutants^44^, in which *Dlx1/2* are selectively deleted in hypothalamic and prethalamic neuroepithelium^12,45,46^. In both global and diencephalic-specific *Dlx1/2* knockouts, the ID domain of Lhx6 expression is absent at E12.5, whereas the TT domain is intact. At E17, we also observe a major reduction in the number of Lhx6-expressing neurons in the ZI (Fig. S14). These results indicate that spatially discrete domains of hypothalamic Lhx6 expression are controlled by the expression of different transcription factors (Fig. S14).

### Nkx2.2-derived Lhx6-expressing neurons in the ZI respond to sleep pressure

Our previous work showed that around 40% of ZI Lhx6-expressing neurons respond to sleep pressure, and ZI Lhx6 neurons promote both REM and NREM sleep ^12^. We sought to identify whether Lhx6 neurons derived from *Nkx2.2*-expressing precursors might selectively respond to sleep pressure. *Nkx2-2* is uniquely expressed in hypothalamic Lhx6 neurons, but *Nkx2-2 is absent* in cortical Lhx6 neurons, unlike *Nkx2-1* and *Dlx1/2*. Our scRNA-Seq analysis and immunostaining indicate that a small number of *Nkx2-2*^*+*^ Lhx6-expressing neurons are located in the postnatal ZI (Fig. S14). In addition, RNA velocity analysis on the combined E12.5 and E15.5 scRNA-Seq datasets indicates that cells in the *Nkx2-2*^+^ cluster in E15.5 are derived from both cells located in the ID and hinge region, indicating potential short-range tangential migration from the hinge region to the ID, which in turn leads to a subset of *Nkx2-2*^*+*^ Lhx6-expressing neurons reaching the ZI (Fig. 7). This is supported by our observation that 28% of Lhx6 ZI neurons express Nkx2.2 at E17.5 (Fig. S12).

**Figure 7.**
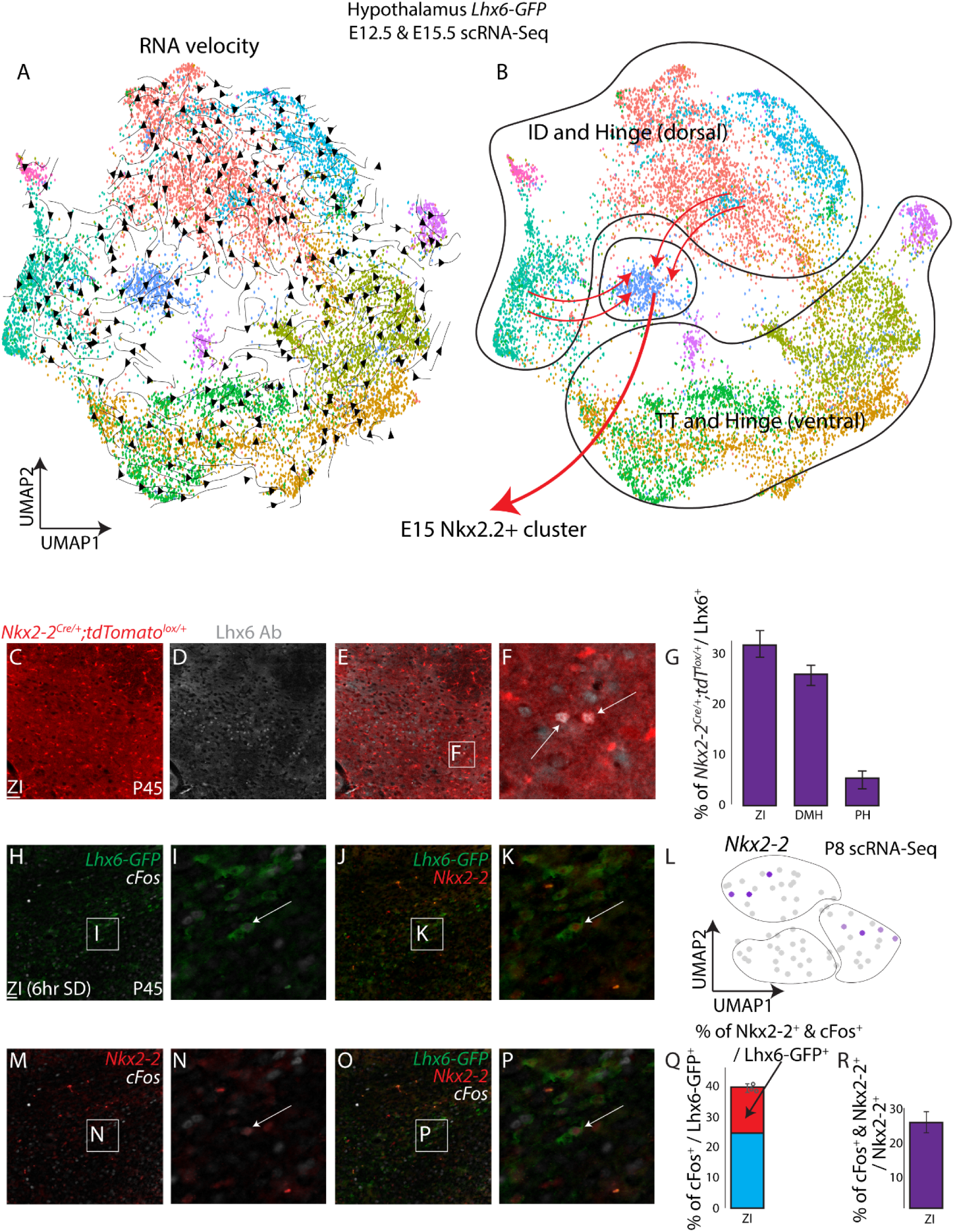
Nkx2.2-expressing Lhx6 ZI neurons respond to increased sleep pressure. (A) UMAP plot with RNA velocity trajectories of E12.5 and E15.5 combined scRNA-Seq dataset. (B) UMAP plot shows distinct domains of hypothalamic Lhx6 neurons. A specific population that continues to express the transcription factor *Nkx2-2* is derived from the ID and dorsal hinge region. (C-F) TdTomato expression in *Nkx2-2*^*Cre/+;*^;*Ai9* mice (red), and Lhx6 antibody staining (grey) identifies Lhx6 neurons in the zona incerta (ZI) are derived from *Nkx2-2*-expressing precursors. (G) A bar graph showing the percentage of tdTomato^+^ and Lhx6-expressing neurons relative to the total number of Lhx6-expressing neurons in the ZI, dorsomedial hypothalamus (DMH, Fig. S15) and posterior hypothalamus (PH, Fig. S15). (H-P) GFP expression from *Lhx6-GFP* (green, H-P), cFos antibody staining (grey) and Nkx2-2 antibody staining (red) shows a specific population of Nkx2-2^+^ Lhx6 neurons in ZI that selectively responds to sleep pressure. (L) UMAP plot showing *Nkx2-2* expression in the anterior portion of Lhx6 neurons. (Q) A bar graph showing the percentage of cFos^+^ and Lhx6-GFP^+^ neurons relative to the total number of Lhx6-GFP^+^ neurons, and demonstrates that a subset of sleep pressure-responsive Lhx6 neurons express Nkx2-2. (R) A bar graph showing the percentage of cFos^+^ and Nkx2-2^+^ neurons relative to the total number of Nkx2-2^+^ neurons, and that a subset of sleep-pressure responding Lhx6 neurons express Nkx2-2. Scale bar = 100 μm.

To determine what fraction of Lhx6-expressing ZI neurons express *Nkx2.2* during development, we performed lineage-tracing analysis using *Nkx2-2*^*Cre/+*^;*Ai9* mice, analyzing the distribution of TdTom/Lhx6-expressing neurons at P45 (Fig. 7, Fig. S14), and observed that ~30% of Lhx6 ZI neurons co-labeled with tdTomato, along with a similar fraction of Lhx6 DMH neurons. In contrast, only a small fraction (~5%) of PH Lhx6-expressing neurons were labeled with tdTomato.

ScRNA-Seq analysis and immunostaining reveal that 30% of Lhx6-expressing ZI neurons continue to express Nkx2.2 in adulthood, raising the question of their potential physiological function. We next then performed 6-hour sleep-deprivation, a robust method to detect cells that respond to sleep pressure, on *Lhx6-GFP* mice^12^ and stained with antibodies to Nkx2-2 and cFos. As shown previously, around 40% of Lhx6-expressing neurons in the ZI responded to sleep pressure, and around 35% of sleep pressure-activated neurons (~15% of all Lhx6-expressing neurons in the ZI) were Nkx2-2^+^ (Fig. 7). In total, 25% of Nkx2-2^+^ ZI neurons express cFos in response to increased sleep pressure. This indicates that *Nkx2.2* may guide the differentiation of a distinct subset of sleep-promoting ZI neurons.

## Discussion

The LIM homeodomain factor Lhx6 is a master regulator of the differentiation and migration of GABAergic neurons of the cortex and hippocampus, as well as many other subcortical telencephalic structures such as striatum and amygdala. Over 70% of cortical interneurons express Lhx6 into adulthood, where it is required for expression of canonical markers of interneuron subtype identity such as Sst and Pvalb^27,32^. In contrast, Lhx6 is expressed in only 1-2% of hypothalamic GABAergic neurons. Lhx6 expression is confined to a broad domain in the dorsolateral hypothalamus, and Lhx6-expressing cells do not undergo widespread long-distance tangential migration. Lhx6-expressing hypothalamic neurons in the ZI play an essential role in promoting sleep^12^, but their function is otherwise uncharacterized. In this study, we seek to characterize the development and molecular identity of hypothalamic Lhx6-expressing neurons, using previous knowledge obtained from studying telencephalic Lhx6-expressing neurons.

In the hypothalamus, in sharp contrast to the telencephalon, Lhx6 is required to prevent neuronal apoptosis (Fig. S16). The fact that loss of function of hypothalamic Lhx6 leads to death of sleep-promoting neurons in the zona incerta may account for the more severe changes in sleep pattern that is seen in the hypothalamic-specific loss of function of Lhx6 than is observed following DREADD-based manipulation of the activity of these neurons^12^. Analysis of *Lhx6*/*Bax* double mutants identified both the Neuregulin and Gdnf signaling pathways as potential neurotrophic mechanisms that promote the survival of hypothalamic Lhx6 neurons. Interestingly, Nrg1/Erbb4-dependent signaling acts as a chemorepellent signal, while Gdnf signaling acts as a chemoattractant, and both regulate the long-range tangential migration of cortical Lhx6 neurons^29,47^. Both signaling pathways may therefore have been at least partially repurposed to regulate cell survival in hypothalamic Lhx6 neurons. The more modest phenotype seen following postnatal loss of function of *Lhx6*, relative to the constitutive mutant, may indicate that the survival of a specific subset of Lhx6-expressing neurons is no longer Lhx6-dependent at later ages.

We observe extensive transcriptional divergence between developing telencephalic and hypothalamic Lhx6 neurons. Notably, we observe clear spatial differences in gene expression among hypothalamic Lhx6 neurons that are not detectable in the MGE. While MGE cells require *Nkx2-1* to activate *Lhx6* expression, *Nkx2-1* is expressed primarily in the TT, in the posterior domain of hypothalamic Lhx6 expression. The TT domain also expresses *Shh* similar to MGE that may regulate *Nkx2-1* expression^18,48^, leading to activation of Lhx6 expression. However, we fail to observe any upstream gene expression (*Shh* or *Nkx2-1*) in MGE scRNA-Seq clusters when the downstream gene is detected (*Nkx2-1* or *Lhx6*)^37^, indicating *Nkx2-1* and *Lhx6* activation could lead to a shutdown of *Shh* and *Nkx2-1* in the MGE. In our hypothalamic Lhx6-expressing neurons, all three genes (*Shh*, *Nkx2-1*, and *Lhx6*) are highly co-expressed in the TT domain, unlike in the MGE.

*Dlx1/2* are expressed in virtually all Lhx6-expressing MGE cells but are not required to maintain *Lhx6* expression^19,21^, while *Dlx1/2* is primarily expressed in the ID domain in the hypothalamus. Furthermore, *Nkx2-2* is not expressed in the telencephalon but is selectively expressed in a previously uncharacterized hinge domain that connects the ID and TT. We find that mutants in *Nkx2-1*, *Nkx2-2*, and *Dlx1/2* selectively eliminate hypothalamic Lhx6 expression in the TT, hinge, and ID domains, respectively. This indicates a high level of spatial patterning and transcriptional diversity among developing hypothalamic Lhx6 neurons. Although hypothalamic Lhx6 neurons do not undergo extensive tangential dispersal, as observed in telencephalon, lineage analysis indicates that by E18, a subset of neurons that express the TT-specific marker Nkx2-1 have migrated to anterior structures such as the ZI. Combined with the observation that Nkx2-2-derived Lhx6 neurons progressively disperse from the hinge domain into the ID implies that subsets of hypothalamic Lhx6 neurons may undergo short-range migration during development.

Lhx6 neurons in the postnatal hypothalamus are likewise highly transcriptionally diverse and do not directly correspond to any of their telencephalic counterparts (Fig. S16). No hypothalamic Lhx6 neurons express *Pvalb*, and only a few selected subsets express either *Sst* or *Npy*. In the cortex, many genes are exclusively expressed in Lhx6-expressing neurons -- including *Sst*, *Pvalb*, and *Npy*. In contrast, in the hypothalamus, no genes were identified that were exclusively expressed in Lhx6 neurons, other than *Lhx6* itself. Neuropeptides such as *Pnoc*, which are expressed in large subsets of hypothalamic Lhx6 neurons, are also widely expressed in many neurons that do not express Lhx6. Finally, molecularly distinct subtypes of Lhx6 neurons are broadly and evenly distributed in the cortex, owing to the widespread tangential dispersal during development. In contrast, in the hypothalamus, we observe clear differences in the expression of neuropeptides and calcium-binding proteins in Lhx6 neurons that broadly correspond to the spatial position of these neurons.

These results provide a starting point to not only better define the molecular mechanisms that control differentiation, survival, and diversification of hypothalamic Lhx6 neurons, but also serve as a molecular toolbox for selectively targeting molecularly distinct neuronal subtypes. Previous studies identified Lhx6 neurons of the zona incerta as being unique in promoting both NREM and REM sleep^12^.

Identification of molecular markers that distinguish different subtypes of Lhx6 neurons in this region can help determine whether this is produced by the activation of distinct neuronal subtypes. We demonstrate that not only are a substantial fraction of Lhx6 ZI neurons derived from *Nkx2.2*-expressing precursors, but that many also continue to express Nkx2.2 into adulthood (Fig. S16). Indeed, Nkx2-2^+^ Lhx6-expressing ZI neurons represent 25% of Lhx6 ZI neurons that express c-fos in response to elevated sleep pressure. Hypothalamic Lhx6 neurons also send and receive connections from many brain regions that regulate innate behaviors, including the amygdala, periaqueductal grey, and ventral tegmental area^12^. The function of these circuits is as yet unknown, and the molecular markers identified in this study can serve as a starting point for investigating their behavioral significance.

## Acknowledgments

This work was supported by a grant from the NIH (R01DK108230) to SB, the Maryland Stem Cell Research Fund (2019-MSCRFF-5124) to DWK, and the Japan Society for the Promotion of Science to TS. We thank Transcriptomics and Deep Sequencing Core (Johns Hopkins) for sequencing of bulk RNA-Seq, bulk ATAC-Seq and scRNA-Seq libraries, and Ross Flow Cytometry Core (Johns Hopkins) for flow sorting of *Lhx6-GFP* cells, and Microscope facility (Johns Hopkins MICFAC, supported by the award number S10OD018118). We thank Marysia Placzek and Wendy Yap for comments on the manuscript.

## Contribution

SB conceived the study. DWK, KL, and SB designed experiments. DWK, KL, ZQW, SZ, AB, MPB, SHL, PWW, and TS performed experiments. DWK, KL, ZQW, SZ, PWW, CS, and TS analyzed data. BL and JLR provided reagents. All authors contributed to writing the paper.

## Conflicts of interest

The authors report no conflicts of interest.

## Data availability

All sequencing data are available on GEO, GSE150687.

## Notes

### Competing Interest Statement

Kai Liu is currently an employee of Genentech. This fact does not, however, affect any of the findings or conclusions of the study.

### Summary of Updates

We have supplemented our scRNA-Seq analysis with RNA velocity (Kallisto with scVelo) to identify developmental trajectories associated with individual Lhx6-expressing regions, and performed regulon analysis (pyScenic) to identify GRN that give rise to diverse Lhx6 neuronal subtypes. We have re-organized the figures to improve overall clarity of the manuscript, and highlighted the potential importance of Nkx2.2 in regulating ZI Lhx6-sleep promoting neurons.

