## Supplemental text and figures for "Gene regulatory networks controlling differentiation, survival, and diversification of hypothalamic Lhx6-expressing GABAergic neurons"

**List of Supplementary Materials**

### Materials and Methods:

#### Mice

All experimental animal procedures were approved by the Johns Hopkins University Institutional Animal Care and Use Committee. All mice were housed in a climate-controlled facility (14-hour dark and 10-hour light cycle) with *ad libitum* access to food and water.

*Lhx6-GFP (Tg(Lhx6-EGFP)BP221Gsat)*[1], *Lhx6*$^{CreER}$ knock-in (B6(Cg)-*Lhx6tm1(*$^{cre/ERT2}$*)Zjh*/J, JAX #010776)[2], *Lhx6*$^{lox/lox}$ [3], *Ai9* (B6.Cg-*Gt(ROSA)26Sortm9(CAG-tdTomato)Hze*/J, JAX #007909)[4], *Bax*$^{lox/lox}$ (B6;129-*Baxtm2Sjk Bak1tm1Thsn*/J, JAX #006329)[5], *Nkx2-2*$^{CreGFP}$ (B6.129S6(Cg)-*Nkx2-2tm4.1(cre/EGFP)Suss*/J, JAX #026880)[6], *Nkx2-1*$^{CreER}$ (*Nkx2-1tm1.1(cre/ERT2)Zjh*/J, JAX #014552)[2], *Foxd1*$^{Cre}$ (B6;129S4-*Foxd1tm1(GFP/cre)Amc*/J, JAX #012463)[7], *Dlx1/2*$^{lox/lox}$ (*Dlx1tm1Rth Dlx2tm1.1Rth*/J, JAX #025612)[8], *Dlx1/2*$^{-/-}$ (gift from John Rubinstein) were used. Mice were time-mated and embryos at various ages (embryonic day (E)11.5, E12.5, E13.5, E15.5, E16.5, E17.5, E18.5, and postnatal day (P) 8) were collected for high-throughput sequencing and histology. Day of birth was considered as P0.

#### Tamoxifen injection

##### *Lhx6*$^{CreER}$ *pulse-chase experiments*

Pups with parental crosses of *Lhx6*$^{CreER/+}$;*Lhx6*$^{lox/+}$;*Bax*$^{lox/+}$;*Ai9* (*Lhx6*$^{CreER/lox}$;*Bax*$^{lox/+}$;*Ai9*) x *Lhx6*$^{CreER/+}$;*Lhx6*$^{lox/+}$;*Bax*$^{lox/+}$;*Ai9* (*Lhx6*$^{CreER/lox}$;*Bax*$^{lox/+}$;*Ai9*) were treated with intraperitoneal 4-Hydroxytamoxifen injection (4-OHT, 0.5 mg/per day, in corn-oil) for 5 consecutive days between P1 and P5. Pups were genotyped on the day of the birth and 3 different genotypes (1. *Lhx6*$^{CreER/+}$;*Ai9*, 2. *Lhx6*$^{CreER/lox}$;*Ai9*, 3. *Lhx6*$^{CreER/lox}$;*Bax*$^{lox/lox}$;*Ai9*) were used. *Lhx6*$^{CreER/CreER}$ genotype dies soon after weaning[2], *Lhx6*$^{CreER/+}$;*Lhx6*$^{lox/lox}$ genotype is not possible to generate due to similar sites of *CreER* and *lox* insertion.

Treated pups were collected between P40 and P45 and processed as described below. Cell counting was conducted in all 3 genotypes in the zona incerta (ZI), dorsomedial hypothalamus (DMH), posterior hypothalamus (PH), S1 somatosensory cortex (CTX), amygdala (AMY) following the Mouse Brain Atlas[9]. Borders were drawn to separate individual regions, using DAPI counterstaining and the Mouse Brain Atlas as a guideline, and 6 500 µm x 500 µm region-of-interest was used to count across cortical layers per section. 3 sections (every second section to avoid counting the same cell) were used per region, and 6 brains that were collected from between 2 and 3 individual litters (different parents) were used. tdTomato expression was observed in blood vessels as previously described[10].

Three different classes of neurons were counted. The first class consists of neurons that only express Lhx6 protein as detected by immunostaining (indicating that no 4-OHT-induced Cre recombination occurred at the *Lhx6* locus). The second class consists of neurons that expressed the only tdTomato but not Lhx6 (indicating Cre-mediated activation of tdTomato, and disruption of *Lhx6*). The third class consists of neurons that expressed both tdTomato and Lhx6 (indicating incomplete 4-OHT-induced Cre recombination, with the induction of tdTomato expression and failure to recombine the conditional allele of *Lhx6*). Only neurons that expressed tdTomato (with or without Lhx6 protein expression) were counted and the total counted the number of neurons used as a denominator. Neurons that only expressed tdTomato were used as a numerator to calculate cell survival rate, as we expect to observe a decrease in the ratio (tdTomato$^{+}$ / (tdTomato$^{+}$ & tdTomato$^{+}$/Lhx6$^{+}$) if *Lhx6* is required for cell survival.

#### ***Nkx2-1$^{CreER}$* pulse-chase experiments**

*Nkx2-1$^{CreER/+}$;Ai9* female mice were time-mated to the same genotype male mice, and 4-OHT was intraperitoneally injected (2 mg) at E11.5, and embryos were collected at E18.5.

#### **Sleep deprivation**

Six-hour sleep-deprivation experiments were performed on *Lhx6-GFP* male mice as previously described[10].

#### **Tissue fixation**

Embryos and mice younger than weaning age (P21) were fixed in 4% paraformaldehyde (PFA) between 8 and 12 hours at 4℃, incubated in 30% sucrose overnight at 4℃, and snap-frozen in OCT compound for histology analysis. Whole embryos were used for fixation until E14.5, and from E14.5, brains were dissected out for fixation. Mice older than weaning age were anesthetized by intraperitoneal injection of avertin and perfused with cold 4% PFA. Brains were post-fixed for 2 hours at 4℃ with 4% PFA and processed as described above.

#### **Cryosectioning**

Frozen brains were sectioned at 25 µm with a cryostat (Leica CM3050S) along either the coronal or sagittal plane, and transferred to Superfrost$^{TM}$ Plus slides.

#### ***In situ* hybridization (ISH)**

Chromogenic and fluorescent *in situ* hybridization was performed as previously described to stain for *Lhx6* (BC065077), *Gfra1* (AW060572), *Gfra2* (BE994145), *Ret* (AW123296), *Dlx1* (BC079609), *Calb1* (AW489595), Calb2 (AI836013), *Gal* (BC044055), *Penk* (AI836252), *Tac1* (BE954293), *Npy* (AI848386), *Sst* (BE984677), *Th* (BF449409), *Gad1* (AW121495), *Nkx2-1* (BC080868), *Nkx2-2* (BG110), *Shh* (BC063087), *Prox1* (BE982394), *Six3* (BE953775), *Lhx8* (BE448496),

*Lef1* (BC038305)[11,12]. RNAscope with probe targeting *Lhx6* was tested on E13.5 mice following the manufacturer's protocol. Images were taken under the Keyence BZ-X800 fluorescence microscope or Zeiss LSM 700 microscope, and processed with ImageJ[13], and pixel density was measured as previously described[14].

#### Immunostaining

Immunostaining was performed with mouse-anti-Lhx6-antibody (1:200, sc-271433, Santa Cruz), rat-anti-RFP (1:500, ABIN334653, antibodies-online), rabbit-anti-Dlx1 (1:500, a gift from Jay Lee), guinea-pig-anti-Dlx1 (1:500, a gift from Jay Lee), rabbit-anti-Nkx2-1 (1:500, EP1584Y, Abcam), mouse-anti-Nkx2-2 (1:100, 74.5A5, DSHB), mouse-anti-NeuN (1:2000, MAB377), rabbit-anti-cFos (1:1000, 226003, Synaptic Systems) as previously described[14], except that M.O.M blocking reagent (MKB-2213) was used following manufacturer's instruction when mouse primary antibodies were used. Alexa Fluor$^{TM}$ 488, 594, 647 secondary antibodies were used in 1:500 dilutions. Sections were mounted with DAPI-Vectamount (Vectorlabs) and imaged under a Keyence BZ-X800 fluorescence microscope or Zeiss LSM 700 microscope. All cell counting was done with ImageJ. Cell counting was conducted in multiple brain areas across developmental ages using standard reference atlases for orientation[11,15], using DAPI counterstaining or NeuN staining as a guideline. For identification of ID and TT, criteria described in our previous study were used[11].  3 sections (every second section to avoid counting the same cell, < E15.5 = 2 sections) were used per region, and 4-6 brains collected from between 2 and 3 individual litters were used. Cell counting was conducted blinded.

#### Bulk RNA-Sequencing

##### *Lhx6 pulse-chase sample sequencing*

*Lhx6$^{CreER/+}$;Ai9* and *Lhx6$^{CreER/lox}$;Bax$^{lox/lox}$;Ai9* P1 pups were treated with 4-OHT as described above and collected at P10. Between 4 and 6 pups from 2 different litters were pooled per sample without regard to sex, and papain-based enzymatic dissociation was performed on the dissected hypothalamus as previously described[16]. Dissociated cells were flow-sorted for tdTomato signal, and between 25,000 and 30,000 cells were collected directly into TRIzol$^{TM}$ LS reagent. RNA was extracted using Direct-zol RNA kits (Zymo Research) and RNA-Sequencing libraries were made using stranded Total RNASeq library prep. 2 libraries were made for *Lhx6$^{CreER/+}$;Ai9*, and 3 libraries were made for *Lhx6$^{CreER/+}$;Lhx6$^{lox/+}$;Bax$^{lox/lox}$;Ai9*. Libraries were sequenced with Illumina NextSeq500, paired-end read of 75 bp, 50 million reads per library. Illumina adapters of sequenced libraries were trimmed using Cutadapt (v1.18)/TrimGalore (v0.5.0)[17] with default parameters, library qualities were assessed using FastQC (v0.11.7)/MultiQC[18]. Libraries were then aligned to mm10 using STAR (v2.54b) [19] with --twopassMode Basic.  RSEM (v1.3.0) was used for quantification[20], with rsem-calculate-expression (--forward-prob 0.5). Expected counts value from RSEM was used to perform differential expression using edgeR (v3.24.3)[21] using default parameters except for calcNormFactors (method = "TMM").

$Lhx6^{CreER/+};Ai9$ or $Lhx6^{CreER/lox};Bax^{lox/lox};Ai9$ enriched genes (fold change > 2 consistent gene value across replicates), were used with EnrichR[22]. $Lhx6^{lox/+};Bax^{lox/lox};Ai9$ enriched genes were compared to the Mouse Cells and Tissues (MESA) dataset available ascot.cs.jhu.edu[23], relying on robustness of expression (NAUC >20) and specificity, as many of the enriched genes detected in this analysis are not strongly expressed in the developing brain.

We reasoned that the genes showing enriched expression in $Lhx6^{CreER/+};Ai9$ relative to $Lhx6^{CreER/lox};Bax^{lox/lox};Ai9$ would be regulated by *Lhx6* and/or *Bax*. Furthermore, since tdTomato expression is detected in blood vessels due to weak Lhx6 expression in endothelial neurons during development[10], we wanted to enrich expression from Lhx6-expressing neurons of the hypothalamus. P8 *Lhx6-GFP*, in which GFP expression is absent in endothelial cells, was used to generate bulk RNA-Sequencing (bulk RNA-Seq) from the cortex and hypothalamus (method described below). Hypothalamus-enriched genes from P8 *Lhx6-GFP* bulk RNA-Seq data were used to enrich genes that are highly expressed in the hypothalamus Lhx6 neurons. After enrichment, the gene lists were compared to single-cell RNA-Sequencing (scRNA-Seq) data from P8 *Lhx6-GFP* hypothalamus using the method described below, to further cross-check specificity of expression and to remove any possible contamination that may occur during flow-sorting from bulk RNA-Seq. EnrichR was used to identify gene pathways, and pathways previously implicated in the regulation of neuronal survival were selected.

***Lhx6-GFP bulk RNA-Seq***

To identify differences between cortical and hypothalamic Lhx6 populations, RNA-Sequencing was performed on E15.5, P0, and P8 *Lhx6-GFP* mice, by collecting 8-10 pups from 2 different litters per library. Libraries were sequenced with Illumina HiSeq 2500, and processed as described in the pipeline described above.

**ATAC-Sequencing**

Cortex and hypothalamus of E15.5 and P0 *Lhx6-GFP* mice were collected, dissociated with papain-based enzymatic reaction, and GFP neurons were flow-sorted. Between 60,000 and 70,000 neurons were collected. Flow-sorted neurons were prepared for ATAC libraries as previously described[24,25]. Libraries were sequenced with Illumina NextSeq500, paired-end read of 75 bp, 50 million reads per library. Each sample was run in duplicate.

Illumina adapters of sequenced libraries were trimmed using Cutadapt (v1.18)/TrimGalore (v0.5.0) and library qualities were assessed using FastQC (v0.11.7)/MultiQC. Libraries were aligned to mm10 using Bowtie 2 (v2.25)[26] using --very-sensitive parameter and Samtools (v1.9)[27] was used to check the percentage of mitochondria DNA reads. Picard (v2.18) was used to remove PCR duplicates, and MACS2 (v2.1.2)[28] was used to capture narrow peaks (open chromatin regions) with --shift 100, --extsize 200, --nolambda, --nomodel parameters. ENCODE blacklist regions of the genome were removed using Bedtools (v2.27) intersect function[28–30]. Bedtools intersect function was used to find matching peaks between replicates, in which the distance between peak ends was less than 10 base pairs. ChIPseeker (v1.18.0)[31] was then used to identify regions that were within 3 kb of the transcription start site (TSS). Footprinting was done using pyDNase (v0.24)[32] wellington_footprints function to find transcription factor binding sites and motif analysis was done on footprinting sites with HOMER (v4.11)[33] with default parameters. Peaks between groups were compared as previously described[24,25] to visualize changes in chromatin accessibility between different ages and brain regions using DiffBind (v.2.10.0) [34] and edgeR using default parameters (FDR < 0.05 & adjusted p-value < 0.05). Differential peaks were compared to bulk RNA-Seq, and open chromatin peaks in promoter regions that correspond to altered gene expression from bulk RNA-Seq were identified to obtain a positive correlation between promoter accessibility and gene expression. Peaks and differential gene expression was then cross-matched to scRNA-Seq, to identify potential different regions within Lhx6 hypothalamic neurons that are demarcated by expression of specific transcription factors.

**Single-cell RNA-Sequencing**

Time-mated E12.5, E15.5, and P8 *Lhx6-GFP* mice were collected, and dissection and dissociation were performed as described previously[16]. Between 6 and 10 embryos/pups from 2 different litters were collected. Following dissociation, $GFP^+$ neurons were flow-sorted using Aria IIu Sorter (BD). Between 20,000 and 25,000 neurons were flow-sorted for E12.5 and E15.5, 2,000 neurons were flow-sorted for P8. Flow-sorted neurons were used for the 10x Genomics Chromium Single Cell System (10x Genomics, CA, USA) using V3.0 chemistry per manufacturer's instruction. Three libraries were sequenced on Illumina NextSeq 500 with ~200 million reads per library. Sequenced files were processed through the CellRanger pipeline (v3.1.0, 10x Genomics) using mm10 genome.

Seurat V3[35] was used to perform downstream analysis following the standard pipeline described previously[36], analyzing neurons that express a high *Lhx6* transcript. Louvain algorithm was used to generate different clusters, and spatial information from individual clusters at E12.5 and E15.5 was identified by referring to our previous hypothalamus scRNA-Seq database HyDD[16], as well as previous analysis of anatomical locations of transcription factors [11]. For P8 scRNA-Seq, region-specific transcription factors that are expressed were compared to E12.5 and E15.5 scRNA-Seq gene lists, as well as matching the identified gene lists to the Allen Brain Atlas ISH data[15]. Previously published scRNA-Seq from E13.5 medial ganglionic eminence (MGE)[37] was processed as described above, and the key markers that label individual clusters were compared to E12.5 Lhx6-expressing hypothalamic neurons.

$Lhx6^+$ neurons across multiple mutant groups ($Foxd1^{Cre/+};Dlx1/2^{lox/lox}$, $Nkx2\text{-}1^{CreER/CreER}$, $Nkx2\text{-}2^{CreGFP/CreGFP}$) from[16], were used to compare the expression level of key transcription factors that define sub-regions of hypothalamic *Lhx6* expression domains.

Previously generated scRNA-Seq datasets from the preoptic region[38], suprachiasmatic nucleus[38,39], ventromedial hypothalamus[40], and whole hypothalamus[16,41–43], were analyzed as described above. GABAergic neurons (*Slc32a1*$^{+}$) were first subsetted from the dataset, and the percentage of neurons expressing *Pnoc*, *Penk*, *Calb1*, *Cck*, *Calb2*, *Gal*, *Tac1*, *Th*, *Npy*, *Trh*, *Sst* was determined.

RNA velocity[44] was used to understand the dynamic state of Lhx6 neuronal development. Kallisto and Bustools[45,46] was used to obtain spliced and unspliced transcripts using --lamanno with GRCm38 mouse genome. Scanpy[47] and scVelo[48] was used to process the Kallisto output with default parameters, based on UMAP coordinates obtained from Seurat.

To identify regulons controlling gene expression in different Lhx6-expressing domains, SCENIC[49] (python implemented pySCENIC (using --masks_dropouts)), was used to calculate regulons using default parameters with mm10 feather files on scRNA-Seq dataset using raw count matrix. Top regulons with z-score higher than 2 were identified as the cluster regulon.

**Statistics**

Two-way ANOVA was used for the Lhx6 pulse-chase experiments in Figure 2 (genotype, brain region). Unpaired t-test was used for all other cell counting studies. The Seurat ‘FindAllMarkers’ function with ‘LR = logistic regression model’ with default parameters was used for analyzing differential gene expression, using the number of total mRNAs and genes as a variable. All bar graphs show mean and standard error of the mean (SEM), with individual data points plotted.

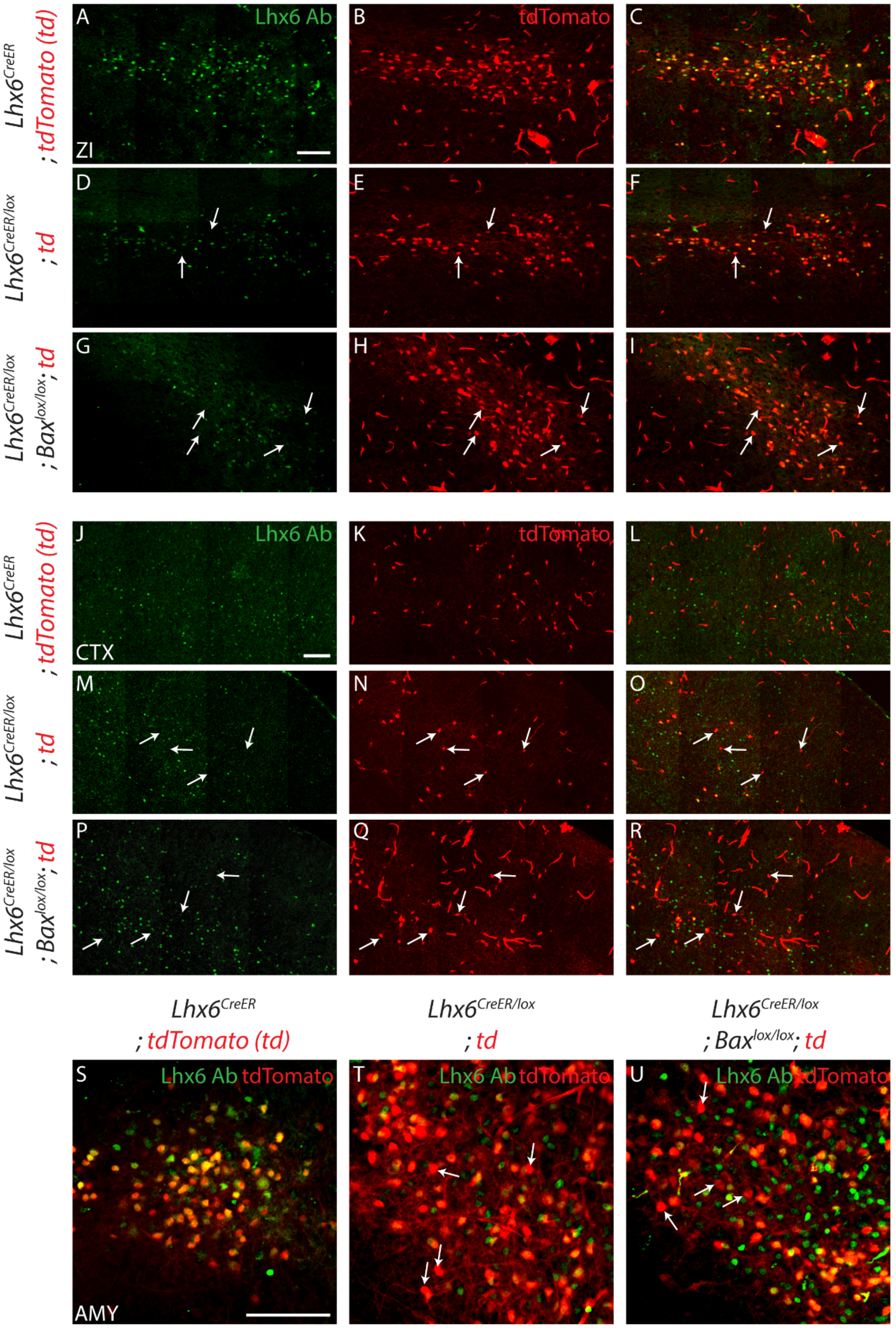


FigS1. Kim, et al.

**Supplemental figure 1.** Representative images of 3 genotypes (1. *Lhx6*$^{CreER/+}$*;Ai9* (A-C, J-L, S), 2. *Lhx6*$^{CreER/+}$*; Lhx6*$^{lox/+}$*;Ai9* (D-F, M-O, T), 3. *Lhx6*$^{CreER/+}$*;Lhx6*$^{lox/+}$*;Bax*$^{lox/lox}$*;Ai9* (G-I, P-R, U) in zona incerta (ZI, A-I), cortex (CTX,

J-R), and amygdala (AMY, S-U) with tdTomato (red) and Lhx6 antibody staining (Lhx6 Ab, green). White indicates tdTomato$^{+}$ neurons without Lhx6 expression. Scale bar = 100 μm. 4-OHT was administered between P1 and P5, and animals were collected between P40 and P45.

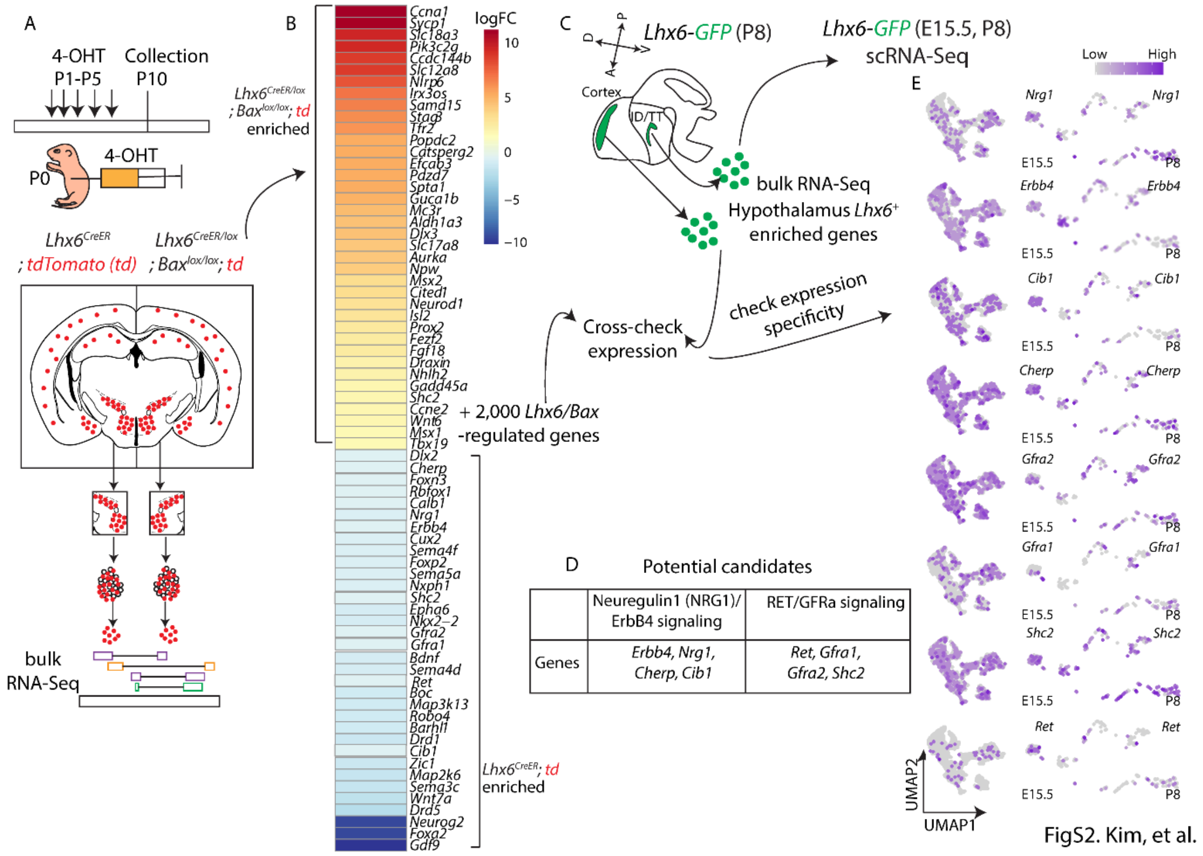


**Supplemental figure 2.** Potential candidates that can regulate survival in hypothalamic Lhx6-expressing neurons. (A) Schematic showing bulk RNA-Seq pipeline from *Lhx6*$^{CreER/+}$*;Ai9* and *Lhx6*$^{CreER/+}$*;Lhx6*$^{lox/+}$*;Bax*$^{lox/lox}$*;Ai9*. (B) A heatmap showing examples of genes (full list in Table. S1) that are enriched in *Lhx6*$^{CreER/+}$*;Ai9* or *Lhx6*$^{CreER/+}$*;Lhx6*$^{lox/+}$*;Bax*$^{lox/lox}$*;Ai9*. Note upregulation of genes that are involved in cell proliferation (*Ccna1*, *Aurka*) and neural precursor neurons (*Irx3os, Cited1, Neurod1*). (C) Schematic showing bulk RNA-Seq from P8 *Lhx6-GFP* cortex and hypothalamus, and scRNA-Seq from E15.5 and P8 *Lhx6-GFP* hypothalamus. (D) Potential candidate genes controlling cell survival can be regulated by *Lhx6* in hypothalamic Lhx6-expressing neurons: Neuregulin-ErbB4 signaling and Gdnf signaling pathways. (E) UMAP plots showing that genes in D are robustly expressed in E15.5 and P8 hypothalamus Lhx6-expressing neurons from the *Lhx6-GFP* scRNA-Seq dataset.

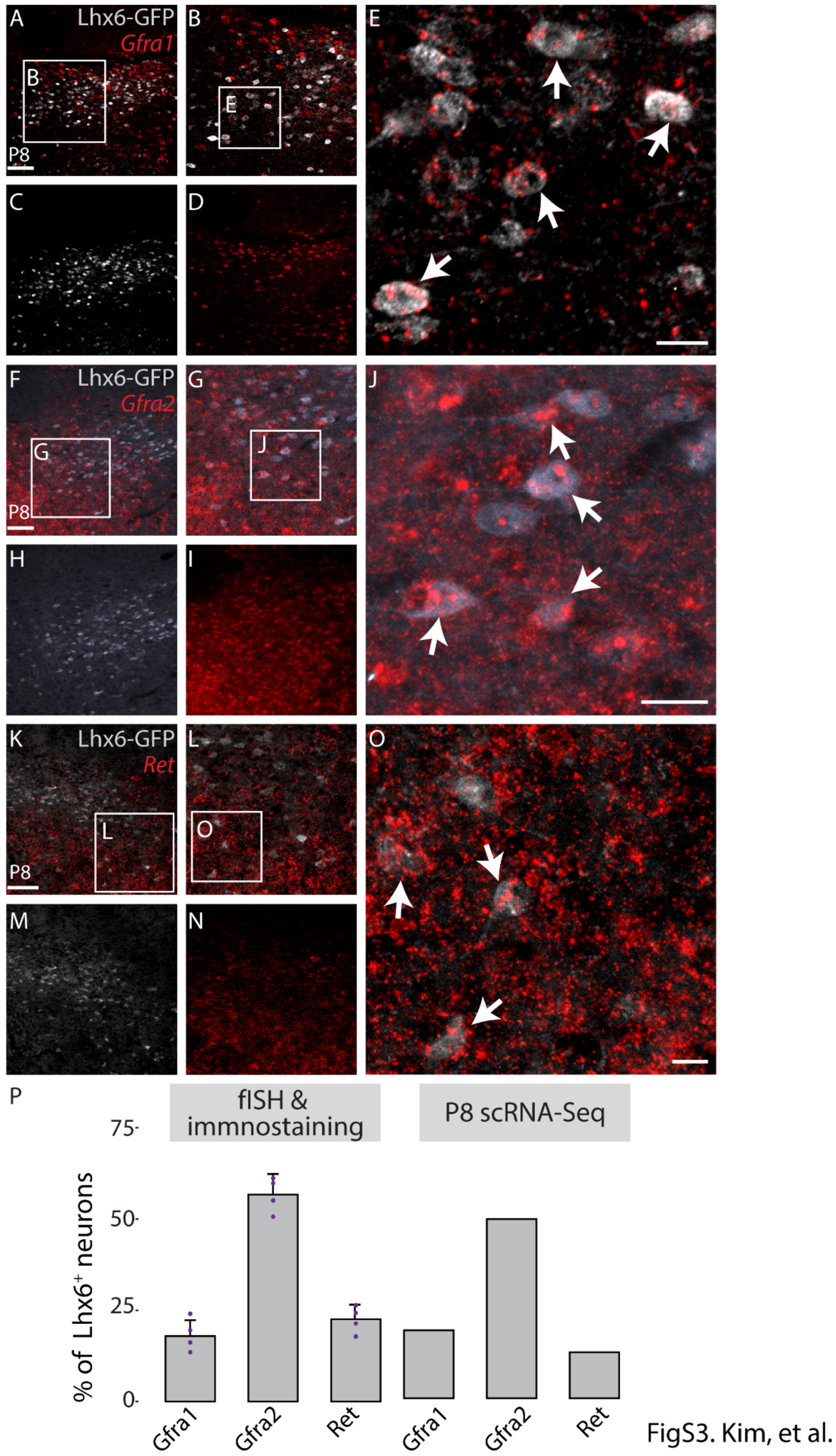


**Supplemental figure 3.** Representative images of *Gfra1* (red, A-E), *Gfra2* (red, F-J), *Ret* (red, K-O) in Lhx6-expressing neurons at P8 in the *Lhx6-GFP* line (grey). White arrows in E, J, O show GFP$^+$ and *Gfra1*$^+$ (E,), *Gfra2*$^+$ (J), *Ret*$^+$ (O). Scale bar = 100 µm (A, F, K), 15 µm (E, J, O). (P) A bar graph showing the percentage of *Gfra1*$^+$/*Gfra2*$^+$/*Ret*$^+$ Lhx6-expressing neurons from 1) fISH (*Gfra1/Gfra2/Ret*, Red) and immunostaining of GFP in *Lhx6-GFP* line (grey) (left), and 2) P8 scRNA-Seq data from flow-sorted hypothalamic GFP$^+$ neurons from *Lhx6-GFP* mice (right).

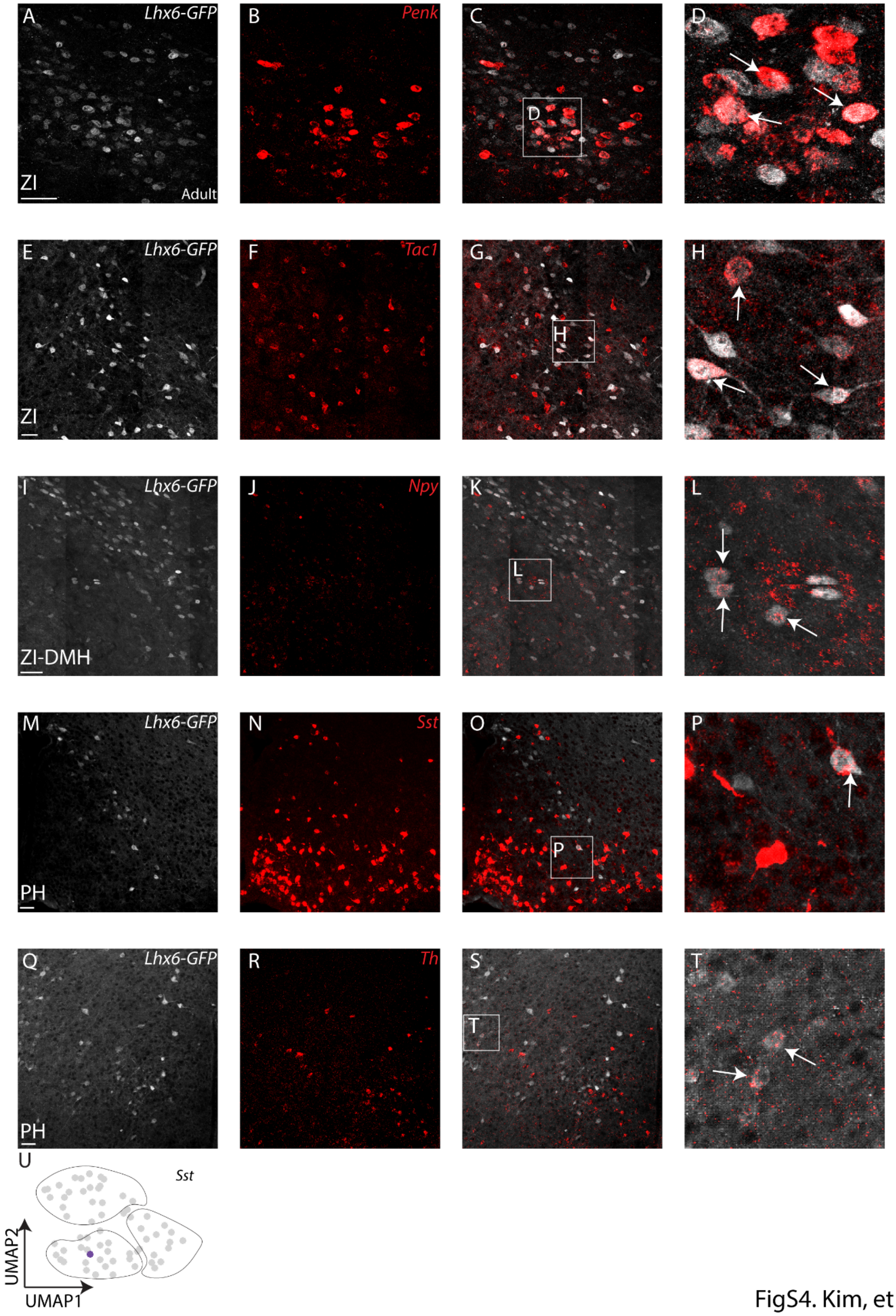


FigS4. Kim, et al.

**Supplemental figure 4.** Fluorescent *in situ* hybridization showing *Lhx6-GFP* (grey) with *Penk* (red, A-D), *Tac1* (red, E-H), *Npy* (red, I-L), *Sst* (red, M-P), and *Th* (red, Q-

T). (U) UMAP plot showing *Sst* expression in ID or TT derived Lhx6-expressing neurons at P8. Scale bar = 50 μm.

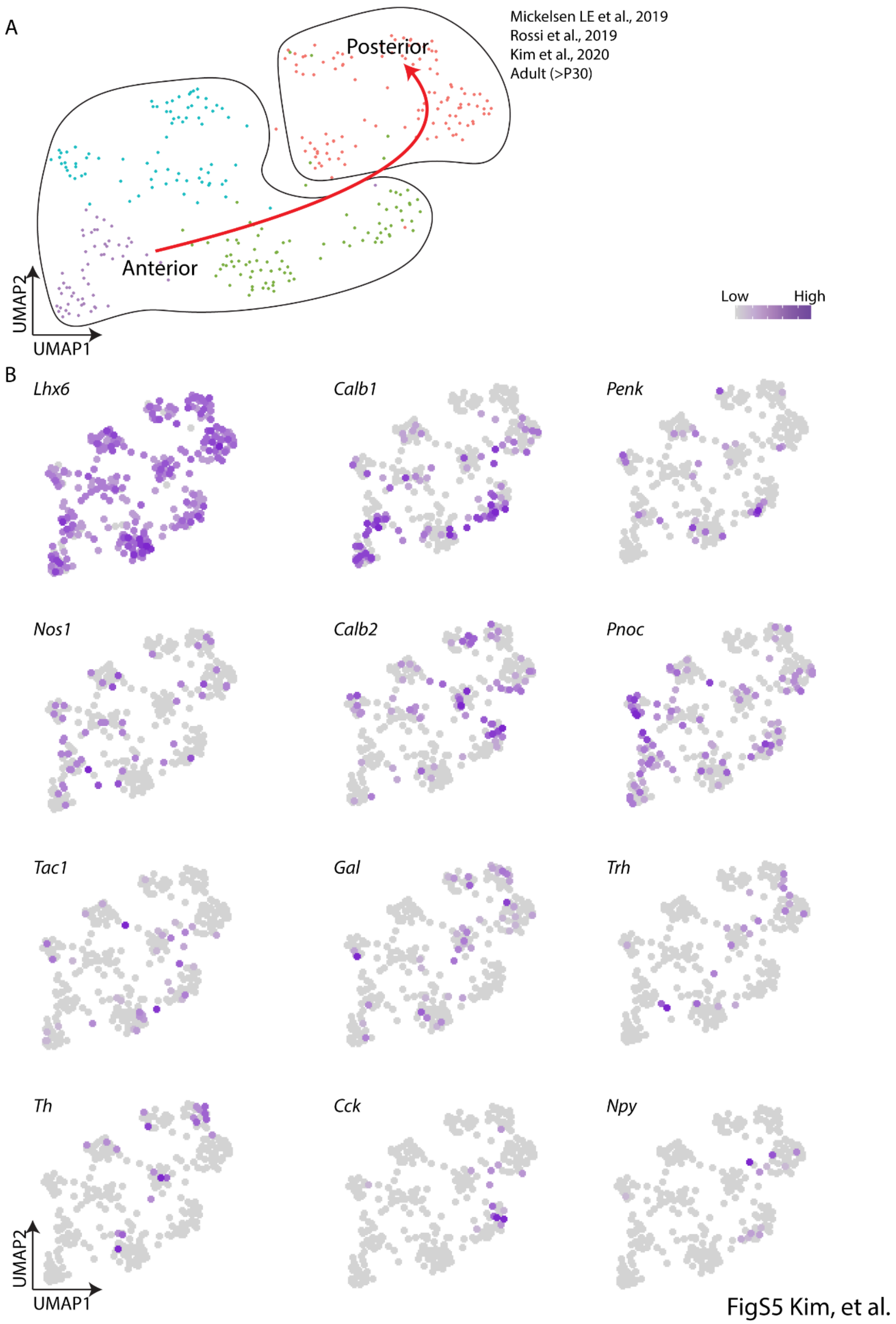


**Supplemental figure 5.** (A) UMAP plot showing Lhx6 neurons in adult (P30>) hypothalamus with the anterior-posterior distribution. (B) UMAP plot showing the expression of neuropeptide and neurotransmitters in adult Lhx6 neurons. Data from [16,50,51].

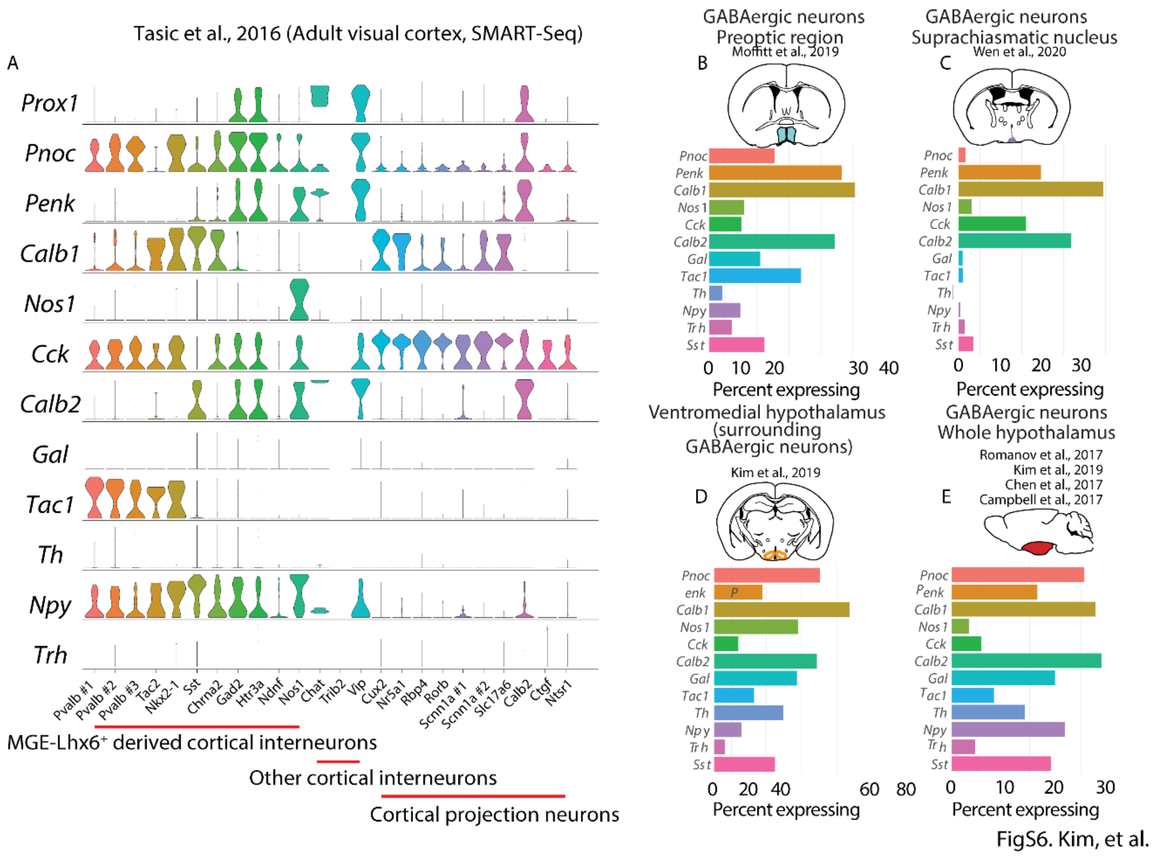


**Supplemental figure 6.** (A) Violin plots showing gene expression in visual cortical neurons of key neuropeptides and transmitters that are expressed in hypothalamic Lhx6-expressing neurons. Data from [52]. (B-E) Neuropeptides and neurotransmitters that are enriched in hypothalamic Lhx6-expressing neurons are widely expressed across GABAergic neurons of the hypothalamus - data from the preoptic area (B)[38], suprachiasmatic nucleus (C)[38,39], ventromedial hypothalamus (D)[40], and whole hypothalamus (E)[16,41–43].

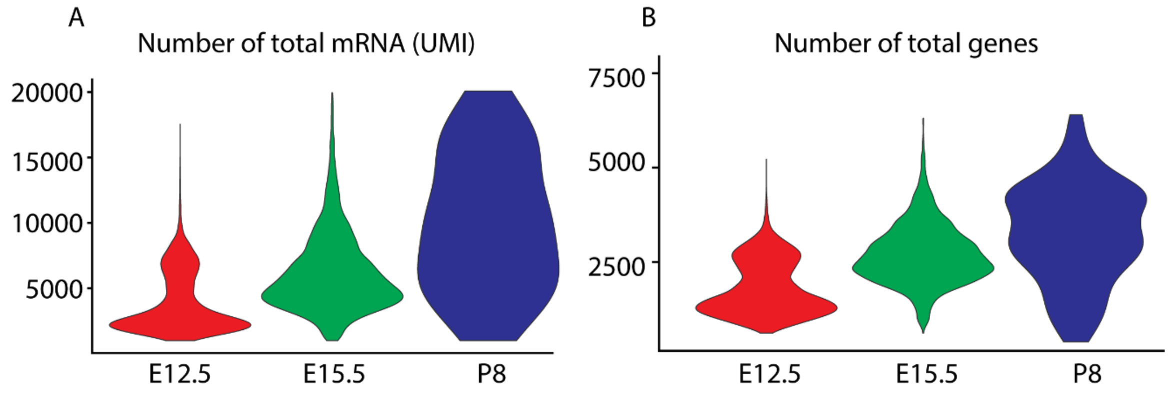


FigS7. Kim, et al.

**Supplemental figure 7.** Violin plots showing the number of total mRNAs (A) and the number of total genes (B) in E12.5, E15.5, and P8 *Lhx6-GFP* scRNA-Seq.

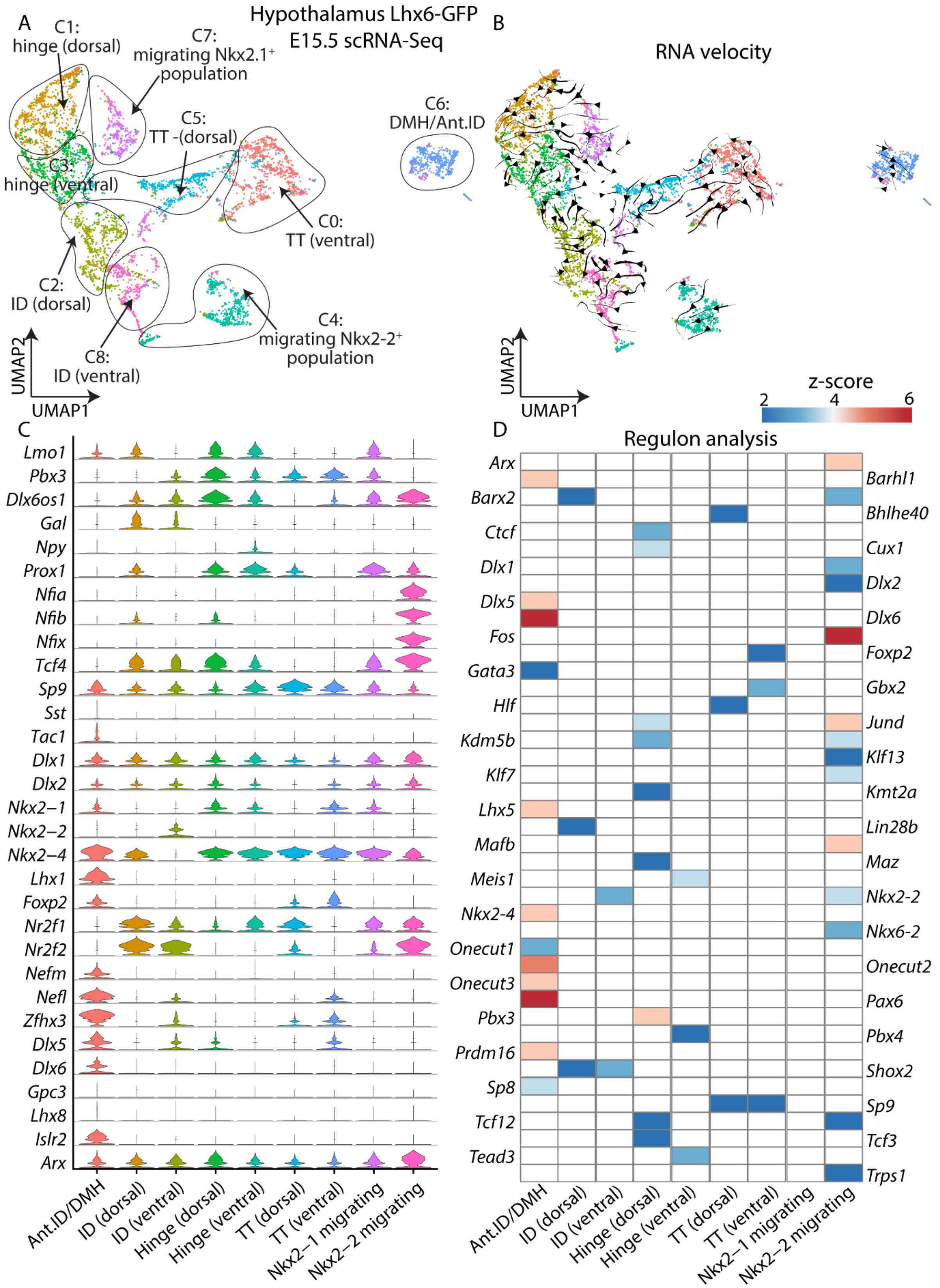


FigS8. Kim, et al.

**Supplemental figure 8.** (A) UMAP plot showing different Lhx6-expressing hypothalamic regions at E15.5. (B) UMAP plot with RNA velocity trajectories. (C) Violin plots showing expression of key transcription factors (and other genes) that are highly expressed in individual domains. (D) A heatmap showing z-scores of significantly differentially expressed regulons between Lhx6-expressing hypothalamic regions. Ant.ID = anterior ID, DMH = dorsomedial hypothalamus.

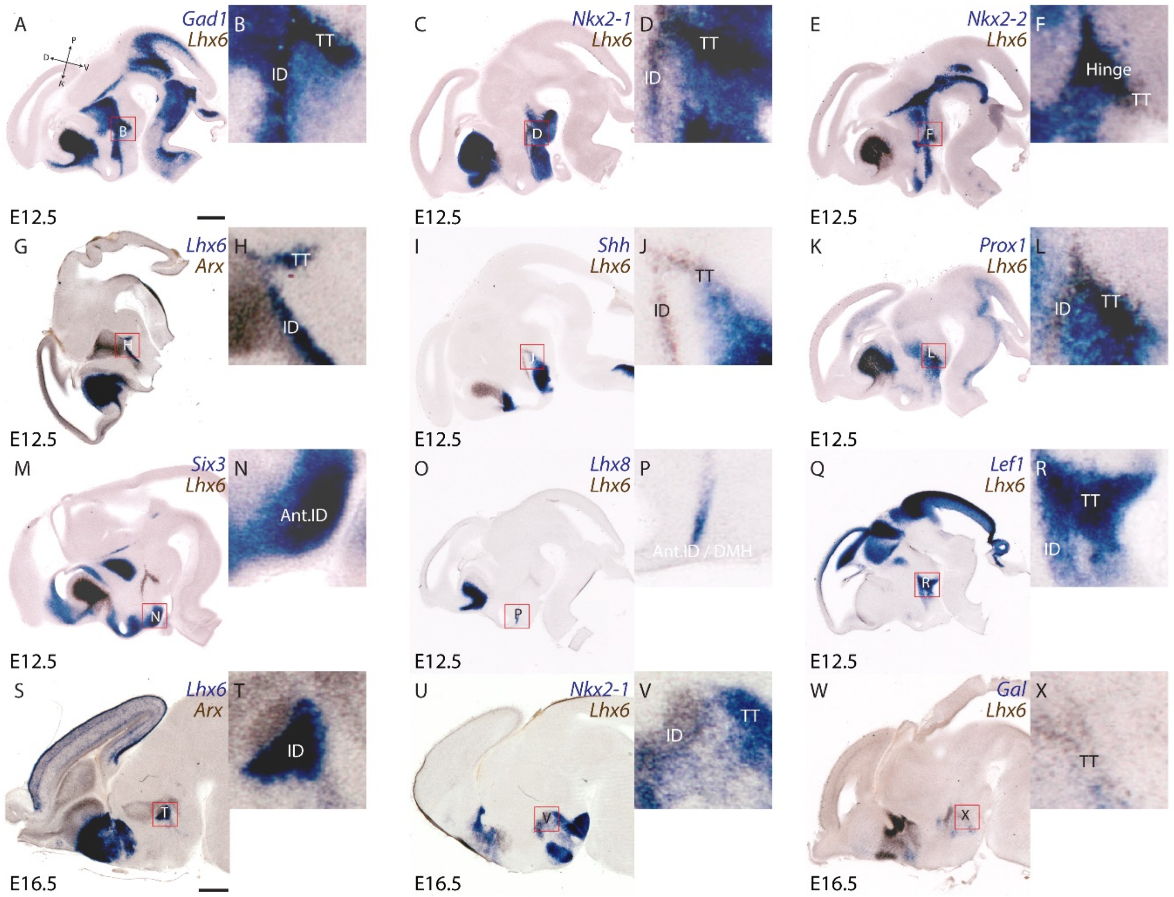


FigS9. Kim, et al.

**Supplemental figure 9**. *In situ* hybridization showing *Lhx6* with *Gad1* (A, B)*, Nkx2-1* (C, D, U, V)*, Nkx2-2* (E, F)*, Arx* (G, H, S, T)*, Shh* (I, J)*, Prox1* (K, L)*, Six3* (M, N)*, Lhx8* (O, P)*, Lef1* (Q, R)*, Gal* (W, X) at E12.5 (A-R) and E16.5 (S-X), shown in sagittal planes. Scale bar = 0.45 mm (A-R), 0.6 mm (S-X) .

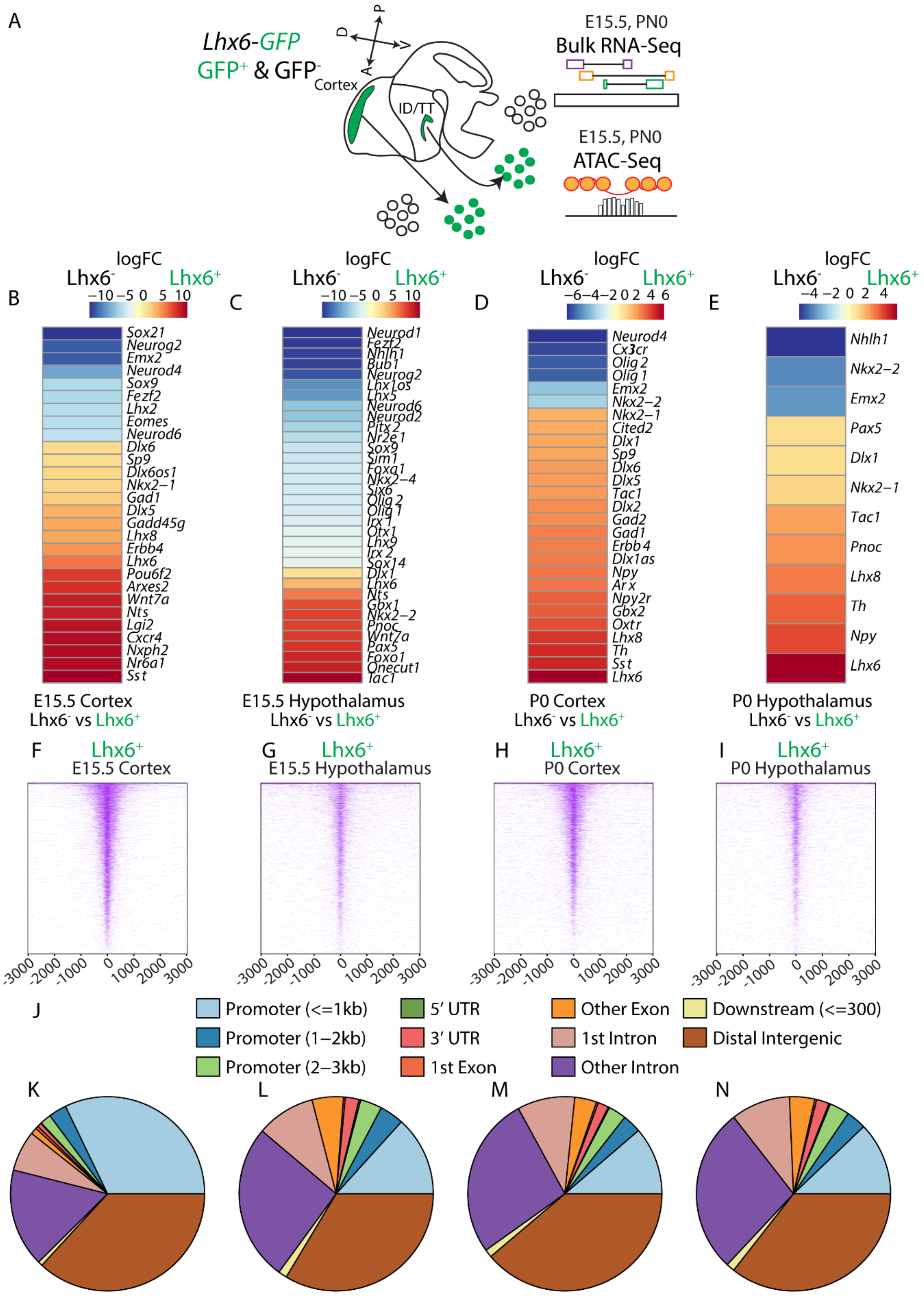


FigS10. Kim, et al.

**Supplemental figure 10.** (A) Schematic showing bulk ATAC-Seq pipeline from flow-sorted *Lhx6-GFP*+ neurons of the cortex and hypothalamus at E15.5 and P0. (B-E) Heatmap showing examples of genes (full list in Table. S2-S3) that are enriched in

*Lhx6-GFP*$^+$ neurons compared to *Lhx6-GFP*$^-$ neurons of the cortex and hypothalamus at E15.5 and P0. (F-I) Heatmap showing open chromatin regions in E15.5 (F, G), P0 (H, I), cortex (F, H), hypothalamus (G, I). (J) Legends for pie graphs in (K-N). (K-N) Distribution of open chromatin regions in E15.5 (K, L), P0 (M, N), cortex (F, H), hypothalamus (G, I).

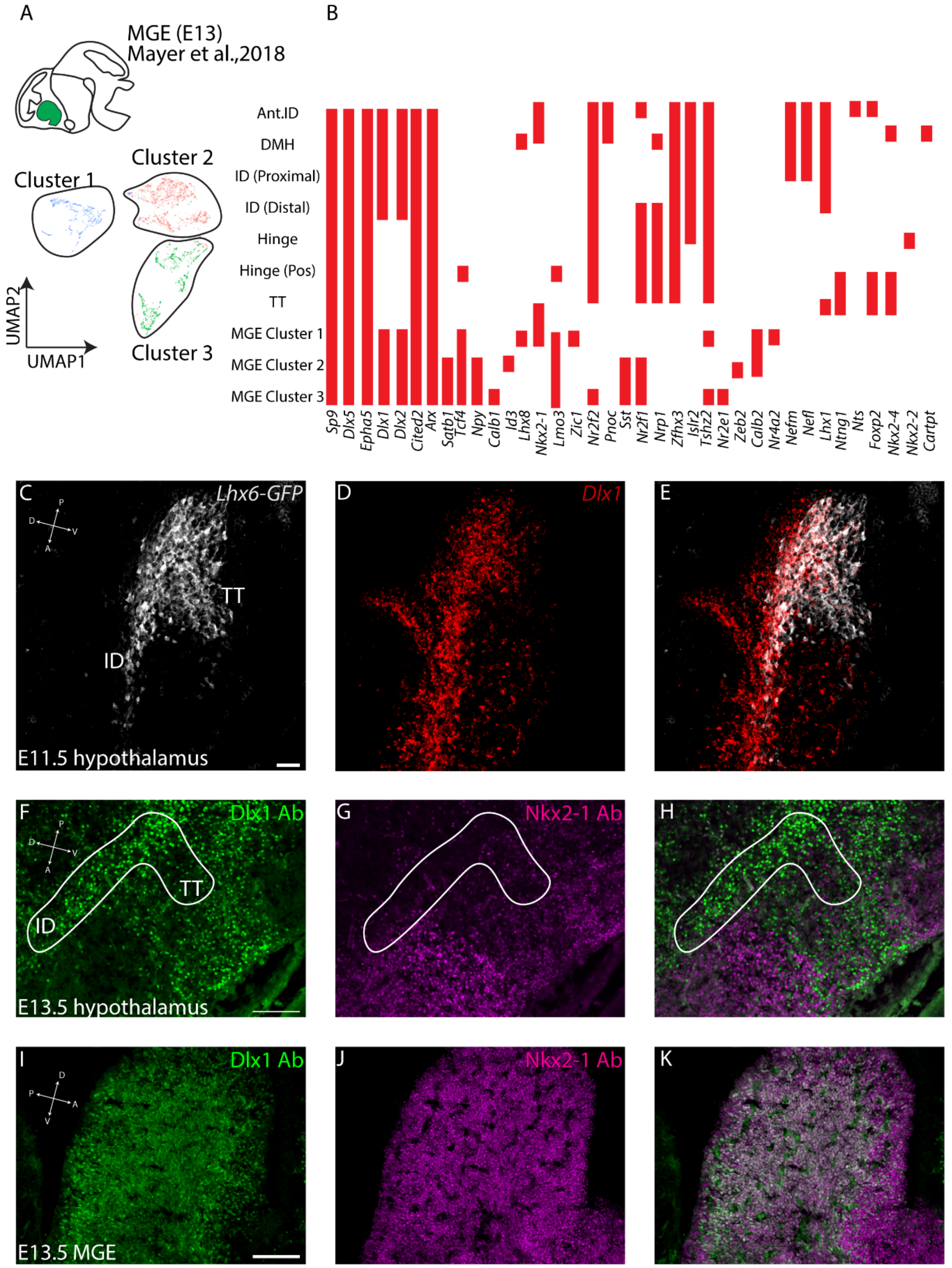


FigS11. Kim, et al.

**Supplemental figure 11.** (A) UMAP plot showing different Lhx6⁺ medial ganglionic eminence (MGE) regions at E13.5. Data from [37]. (B) Graphs showing presence (red bars) of key genes that are expressed in hypothalamic and/or MGE LHx6⁺

populations. Note a lack of overlap in gene expression profiles between hypothalamus MGE $Lhx6^{+.}$ populations. (C-E) fISH showing GFP expression in *Lhx6-GFP* line at E11.5 (grey) and *Dlx1* (red). Note *Dlx1* covers most of the ID. (F-H) Immunostaining of Dlx1 (green) and Nkx2-1 (magenta) in E13 ID and TT. Note the lack of co-expression Dlx1 and Nkx2-1, and Dlx1/2 and Nkx2-1 expression delineate separate zones in the ID and TT.  (I-K) Immunostaining of Dlx1 (green) and Nkx2-1 (magenta) in E13 MGE. Note a high level of co-expression Dlx1 and Nkx2-1. Scale bar = 50 μm.

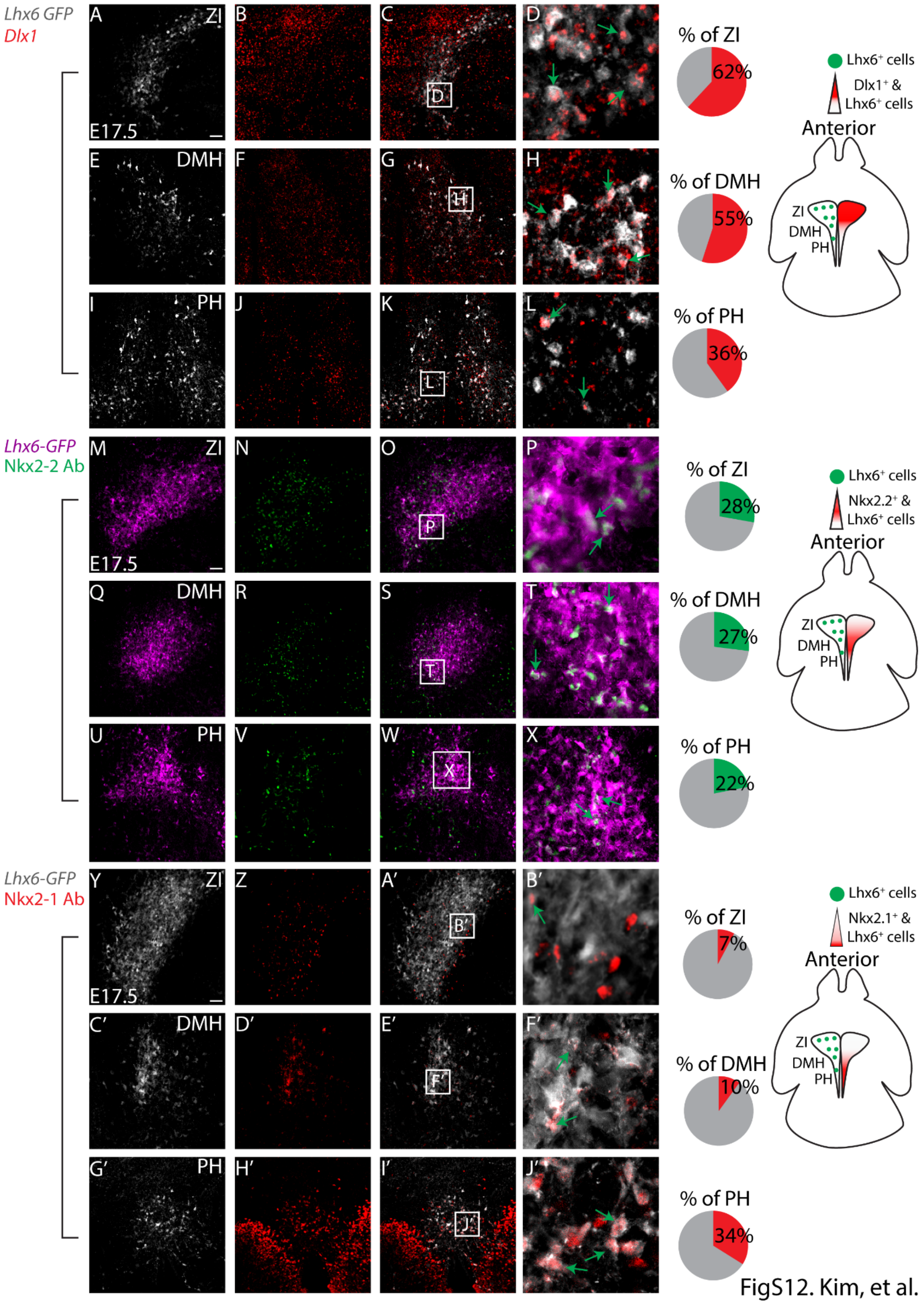


**Supplemental figure 12.** Staining showing distribution and the percentage of *Dlx1* (A-L *Lhx6-GFP* in grey, *Dlx1* in red), *Nkx2-2* (M-X, *Lhx6-GFP* in magenta, Nkx2-2 in green), and *Nkx2-1* (Y-J’, *Lhx6-GFP* in grey, Nkx2-1 in red) in E17.5 zona incerta (ZI,

A-D, M-P, Y-B’), dorsomedial hypothalamus (DMH, E-H, Q-T, C’-F’), and posterior hypothalamus (PH, I-L, U-X, G’-J’). Scale bar = 50 μm.

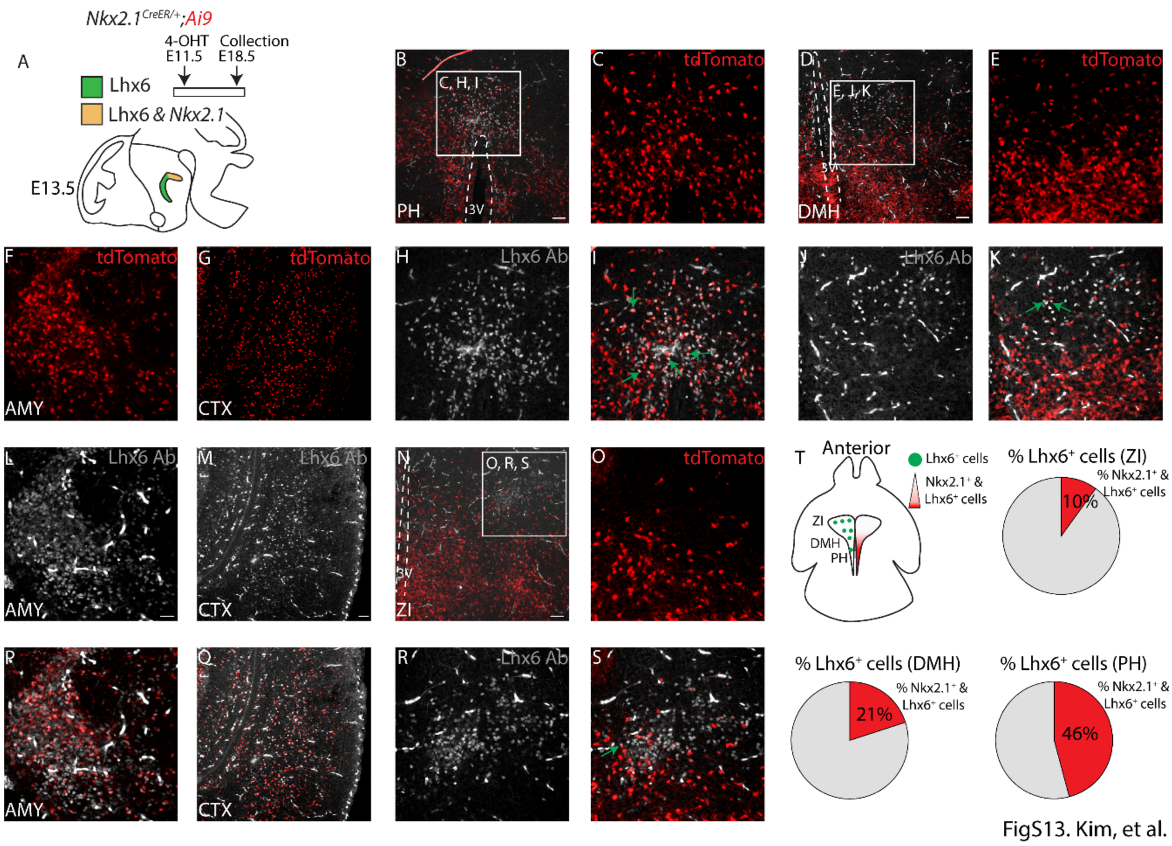


FigS13. Kim, et al.

**Supplemental figure 13.** Nkx2-1 is required only for the specification of posterior hypothalamic Lhx6-expressing neurons. (A) Schematic showing 4-OHT treatment into E11 *Nkx2-1*$^{CreER/+;}$*Ai9* dams and collection of embryos at E18.5 (top), and schematic showing distribution of *Nkx2-1*$^{-}$*/Lhx6*$^{+}$ in ID (anterior) and *Nkx2-1*$^{+}$*/Lhx6*$^{+}$ in TT (posterior). (B-S) Immunostaining showing Lhx6 (grey) and tdTomato (red, *Nkx2-1*$^{CreER/+;}$*Ai9*) in the amygdala (AMY, F, L, P), cortex (CTX, G, M, Q), posterior hypothalamus (PH, B, C, H, I), dorsomedial hypothalamus (DMH, D, E, J, K), and zona incerta (ZI, N, O, R, ST). Green arrows show co-localization. (T) Schematic showing horizontal mouse brain section highlighting ZI, DMH, PH, and distribution of hypothalamic Lhx6-expressing neurons and Nkx2-1$^{+}$/Lhx6$^{+}$ neurons are shown (top left). Pie graphs showing the percentage of Nkx2-1$^{+}$/Lhx6$^{+}$ neurons. Note a posterior bias in the distribution of Nkx2-1$^{+}$/Lhx6$^{+}$ neurons. Scale bar = 50 µm.

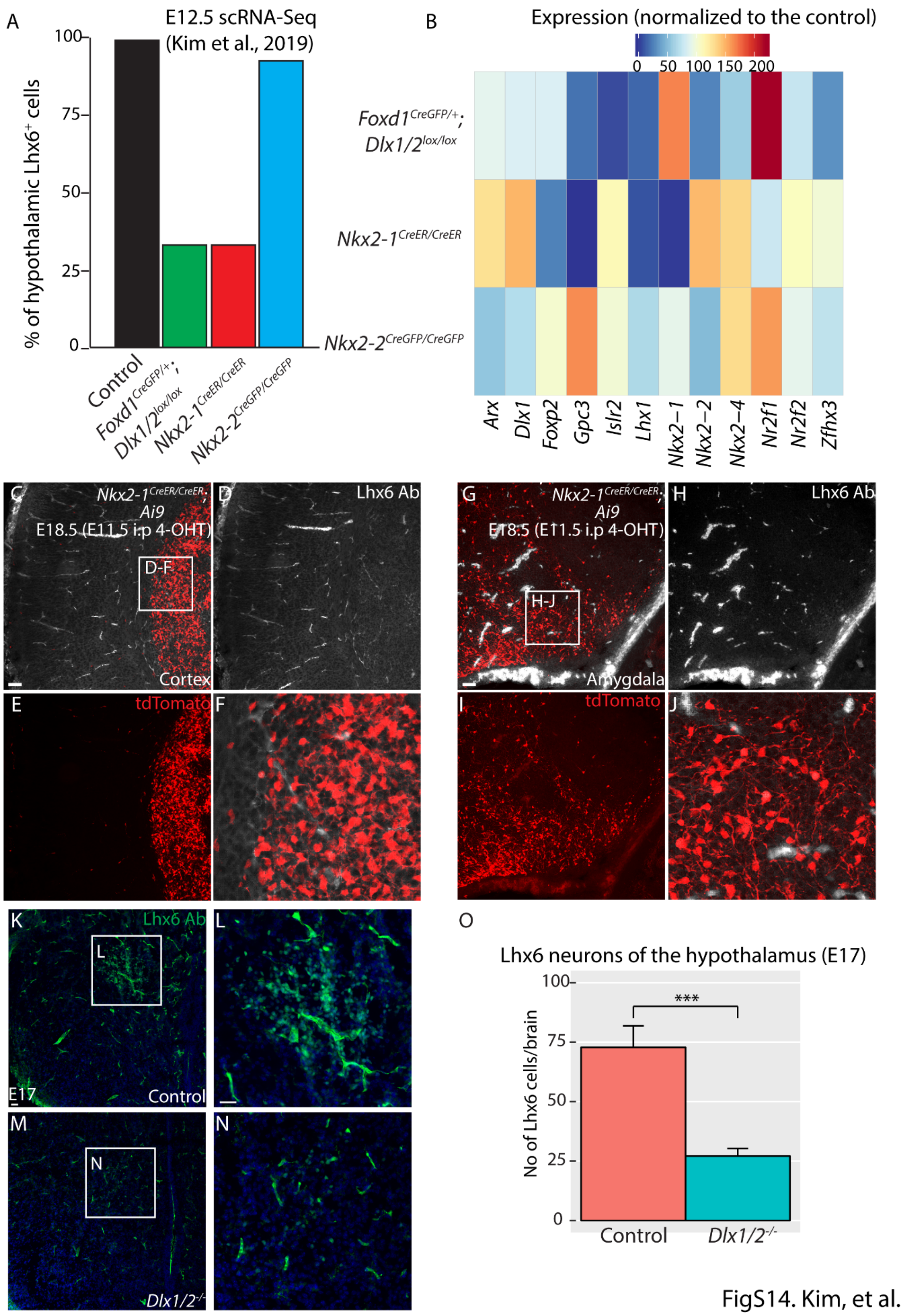


**Supplemental figure 14.** (A-B) scRNA-Seq result from [16], *Foxd1*$^{Cre/+}$;*Dlx1/2*$^{lox/lox}$, *Nkx2-1*$^{CreER/CreER}$, *Nkx2-2*$^{CreGFP/CreGFP}$ from hypothalamic *Lhx6*-expressing neurons. (A) Bar graphs showing the percentage of neurons that express *Lhx6* across 4

genotypes. (B) Heatmap showing changes in the percentage of cells expressing individual transcription factors across 4 genotypes. (C-J) *Nkx2-1*$^{CreER/CreER;}$*Ai9* (4-OHT treatment at E11.5, collection at E18.5) showing Lhx6 antibody staining (grey) and tdTomato (red) in the cortex (C-F) and amygdala (G-J). Note the absence of Lhx6-expressing neurons in both brain regions. Scale bar = 50 µm. (K-O) Lhx6 expression (green) in control (K, L) and *Dlx1/2*$^{-/-}$ (M, N) at E17.5 ZI. The number of Lhx6-expressing neurons is shown in (O). ZI = zona incerta. Scale bar = 50 µm. *** $p < 0.05$

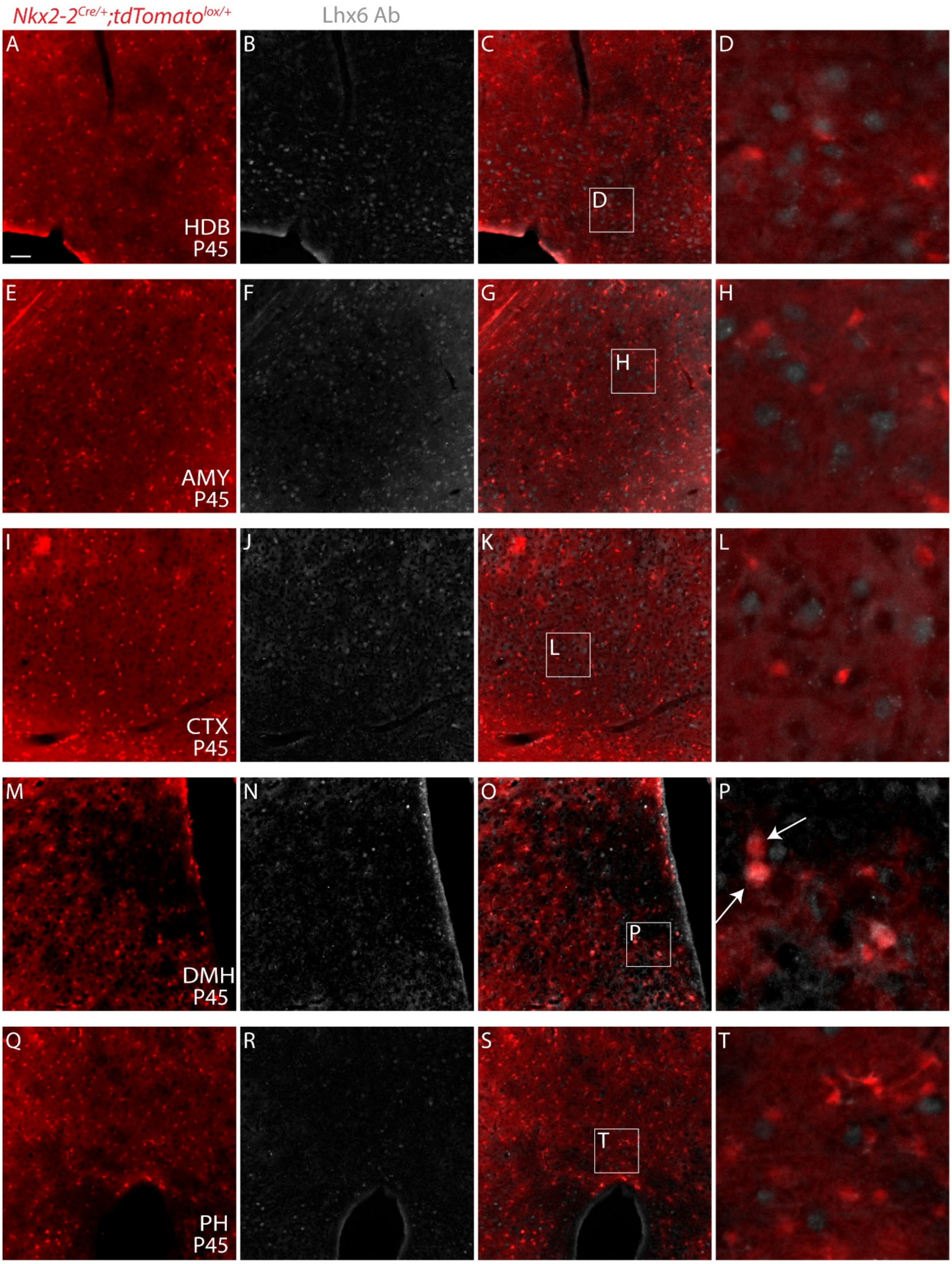


FigS15. Kim, et al.

**Supplemental figure 15.** tdTomato expression from *Lhx6*$^{Cre/+;}$*;Ai9* line (red), and Lhx6 antibody staining (grey) in the diagonal band of Broca (HDB, A-D), amygdala

(AMY, E-H), cortex (CTX, I-L), dorsomedial hypothalamus (DMH, M-P), posterior hypothalamus (PH, Q-T).

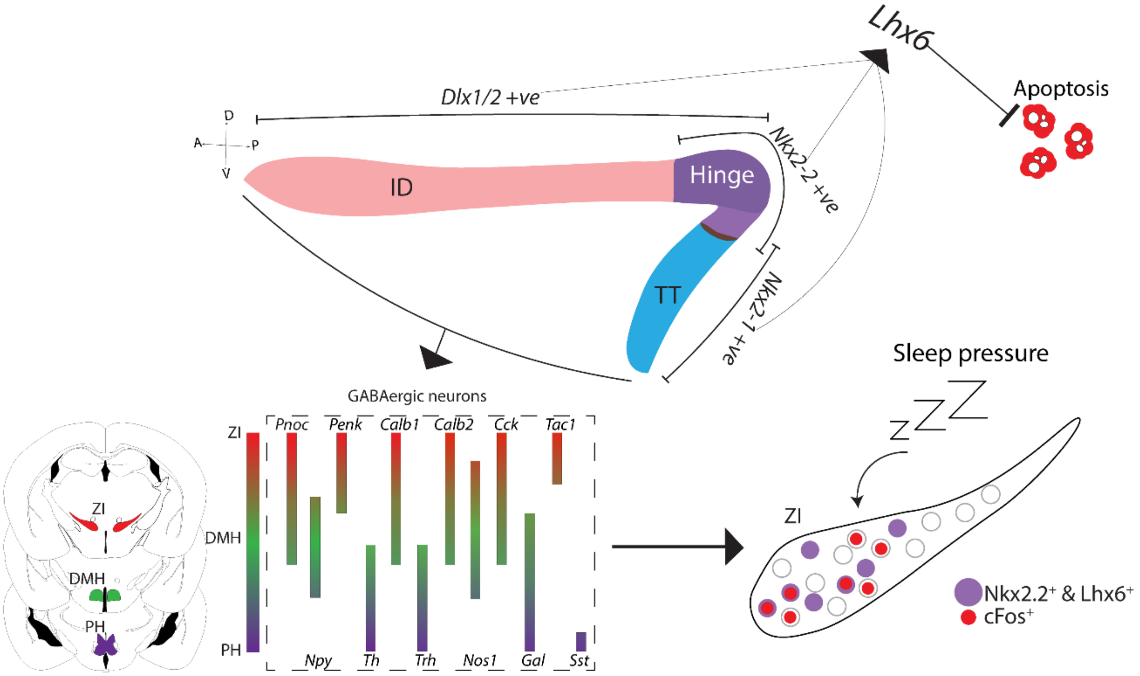


FigS16. Kim, et al.

**Supplemental figure 16.** Schematic summary of hypothalamic Lhx6 development and diversity.

**Table S1.** Differential gene lists from bulk RNA-Seq between *Lhx6*$^{CreER/+}$*;Ai9* and *Lhx6*$^{CreER/+}$*;Lhx6*$^{lox/+}$*;Bax*$^{lox/lox}$*;Ai9***.**

**Table S2.** Differential gene expression in P8 hypothalamic Lhx6 scRNA-Seq data.

**Table S3.** Differential gene expression in E12.5 hypothalamic Lhx6 scRNA-Seq data.

**Table S4.** Differential gene expression in E15.5 hypothalamic Lhx6 scRNA-Seq data.

**Table S5.** Differential gene lists from bulk RNA-Seq between cortical and hypothalamic *Lhx6-GFP*$^{+}$ neurons at E15.5 and P0.

**Table S6.** Differential gene lists from bulk RNA-Seq between *Lhx6-GFP*$^{+}$ and *Lhx6-GFP*$^{-}$ neurons of cortex and hypothalamus at E15.5 and P0.

**Table S7.** Differential peaks from ATAC-Seq between cortical and hypothalamic *Lhx6-GFP*$^{+}$ neurons at E15.5.

**Table S8.** Differential peaks from ATAC-Seq between cortical and hypothalamic *Lhx6-GFP*$^{+}$ neurons at P0.

**Table S9.** Differential gene expression in E13 MGE Lhx6 scRNA-Seq data. Data from [37].
